# Key-cutting machine: A novel optimization framework for tailored protein and peptide design

**DOI:** 10.1101/2025.01.05.631393

**Authors:** Yan C. Leyva, Marcelo D. T. Torres, Carlos A. Oliva, Cesar de la Fuente-Nunez, Carlos A. Brizuela

## Abstract

Computational protein and peptide design is emerging as a transformative framework for engineering macromolecules with precise structures and functions, offering innovative solutions in medicine, biotechnology, and materials science. However, current methods predominantly rely on generative models, which are expensive to train and inflexible to modify. Here, we introduce the Key-Cutting Machine (KCM), a novel optimization-based platform that iteratively leverages structure prediction to match desired backbone geometries. KCM requires only a single GPU and enables seamless incorporation of user-defined requirements into the objective function, circumventing the high retraining costs typical of generative models while allowing straightforward assessment of measurable properties. By employing an Estimation of Distribution Algorithm, KCM optimizes sequences based on geometric, physicochemical, and energetic criteria. We benchmarked its performance on α-helices, β-sheets, and unstructured regions, demonstrating precise backbone geometry design. As a proof of concept, we applied KCM to antimicrobial peptide (AMP) design by using a template AMP as the “key”, yielding a candidate with potent *in vitro* activity against multiple bacterial strains and efficacy in a murine infection model. KCM thus emerges as a robust tool for *de novo* protein and peptide design, offering a flexible paradigm for replicating and extending the structure–function relationships of existing templates.

## Introduction

Protein and peptide design aims to construct synthetic proteins and peptides or modify existing ones to achieve new functionalities^1,2^. Successful designs include proteins that inhibit viral infections such as influenza^3,4^, SARS-CoV-2^5^ and HIV^6^, as well as enzymes that catalyze reactions for which no natural counterparts were previously known^7^. Additional applications include biosensors to detect various molecules^8^, including fentanyl^9^, self-assembling protein nanoparticle vaccines for SARS-CoV-2^10^, self-assembling nanomaterials^11^ and biological logic gates^12^. In the case of peptides, a simple generative model was proposed to design new antimicrobial peptides against *Cutibacterium acnes*^13^. ProT-Diff^14^, another generative model, produced novel peptides; of these, 34 showed antibacterial activity against both Gram-positive and Gram-negative bacteria, as well as *in vivo* activity against a clinically relevant drug-resistant *E. coli* strain. These advances highlight the broad therapeutic and biotechnological potential of computational protein and peptide design^15^.

Computational protein and peptide design is often framed as a combinatorial optimization problem, where the goal is to identify an amino acid sequence that folds into a structure with minimal free energy^1^. This challenge is partially mitigated by discretizing side-chain conformations (rotamers)^16^. Thus, the task reduces to finding the optimal combination of rotamers that minimize free energy for a given backbone. Alternative formulations include mixed-integer linear programming^17^ and cost function networks^18^. However, the protein design problem remains extraordinarily complex^19,20^.

Deep learning methods have emerged as powerful tools for protein design. Notable examples include ProteinMPNN^21^, a message-passing neural network; ProteinSolver^22^, a graph-based neural network; and ESM-IF1^23^, which integrates Geometric Vector Perceptron (GVP) with a Transformer. Given a structure in PDB format, these models predict the amino acid sequences most likely to fold into the provided structure. Additional improvements can be made by incorporating extra information, as shown in SPDesign^24^, SPIN-CGNN^25^, and PiFold^26^.

Recently, generative models, including diffusion models, have been applied to protein design. These models, which include RFDiffusion^27^ and GRADE-IF^28^, can generate novel backbone structures from random noise that can then be paired with protein design networks (e.g., ProteinMPNN) to produce corresponding sequences. Additionally, language models like StructureGPT^29^ or Tpgen^30^ have shown promise in protein design.

However, a major limitation of existing generative models is their substantial computational cost, primarily due to the need for retraining from scratch whenever a new measurable property is introduced into the loss function. This retraining procedure requires significant computational resources, rendering iterative design tasks impractical. To overcome this limitation, we propose a novel optimization-based model for protein and peptide design—referred to as the Key-Cutting Machine (KCM) model (**Figure 1a**). KCM requires only a single GPU and enables the seamless incorporation of user-defined requirements into the objective function, thus avoiding the repeated retraining overhead typical of generative models and simplifying the assessment of measurable properties.

**Figure 1.**
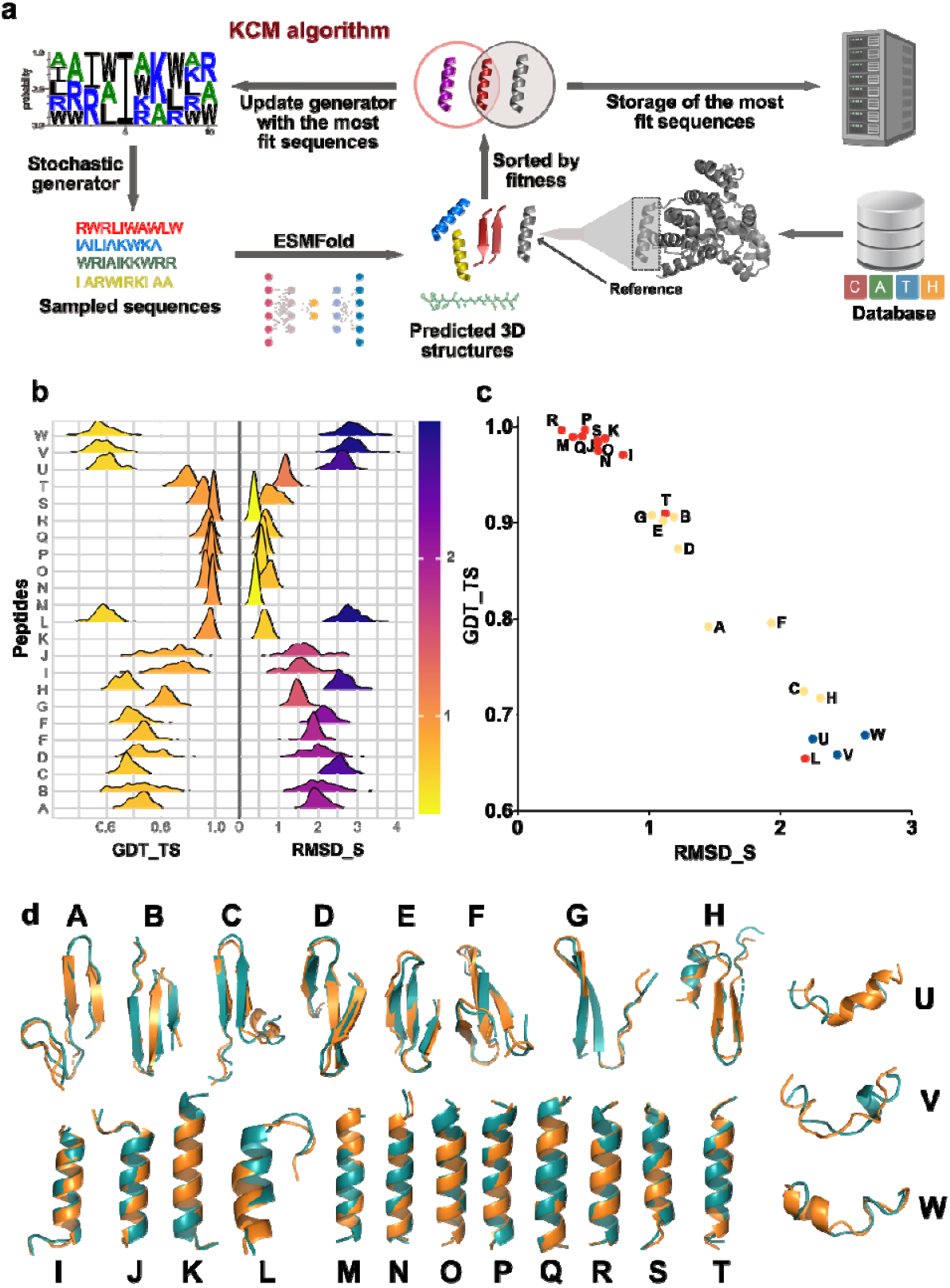
Key-cutting machine algorithm. **(a)** Schematic of the KCM algorithm including the stochastic generator and the three-dimensional structure predictor (ESMFold). **(b)** Distribution for GDT_TS and RMSD_S from the best 100 designs according to their fitness, for each protein. Continuous color range for both, GDT_TS and RMSD_S, with colors going from yellow (zero) to purple (four). **(c)** GDT_TS as a function of RMSD_S of the highest fitness design for each protein. Mainly β-sheets in yellow, α-helix in red and those without defined secondary structure in blue. **(d)** Backbone superposition of the reference protein (green) with respect to the backbone of the highest fitness design (orange). Correspondence between characters and proteins for **(b), (c)**, and **(d)**: A-1EMN, B-1LQL, C-1NP6, D-1O6W, E-2E45, F-2KXQ, G-1S7M, H-1P9N, I-5UIY, J-5UIU, K-3SB1, L-2QQ8, M-3M9Q, N-3H25, O-3EWK, P-3C8V, Q-2QIW, R-2OAR, S-2LKM, T-1MSL, U-3W68, V-1R5L, W-1OIP.

In this framework, the “key” corresponds to the target structure and its sequence, and a structure-prediction method generates multiple copies of this key. KCM then refines these sequences by comparing the original key with its predicted copies. We implement an estimation of distribution algorithm (EDA)^31^ using an island model and employ ESMFold as the structure predictor to guide the optimization process.

As a proof-of-concept, we chose a 12-residue peptide with antimicrobial properties^32^. We computationally designed derivatives of this small peptide, synthesized them chemically, and tested their antimicrobial activity both *in vitro* and *in vivo*.

## Results

Our methodology consists of three stages (**Figure 1**). First, we define an optimization model for the KCM approach, characterized by an objective function to be maximized. Next, we propose an EDA to solve this model. Finally, we apply the algorithm to a dataset of proteins with known sequences and secondary structures, including α-helices, β-sheets, and unstructured proteins.

The protein design problem is daunting because of the immense sequence space and the unpredictable mapping from amino acid sequences to structures^33^. Even a single amino acid mutation can dramatically alter the structure of a given protein or peptide. We hypothesize that our EDA can accurately estimate the distribution of structures within sequence space^34,35^.

### Protein Design: Three types of secondary structures

In our experiments, protein designs dominated by α-helices required fewer generations to converge than their β-sheet counterparts, in part because α-helices are typically shorter. A termination condition of 100 generations was applied to α-helical proteins (e.g., PDBs: 5UIY, 5UIU, 3SB1, 2QQ8, 3M9Q, 3H25, 3EWK, 3C8V, 2QIW, 2OAR, 2LKM, and 1MSL) and proteins lacking defined secondary structure (PDBs: 3W68, 1R5L, and 1OIP). For β-sheet proteins (PDBs: 1EMN, 1LQL, 1NP6, 1O6W, 2E45, 2KXQ, 1S7M, 1P9N), we used a termination condition of 1,000 generations or until the Global Distance Test Total Score (GDT_TS)^36^ reached at least 0.9. When the threshold was met earlier (e.g., PDBs: 1O6W, 2E45, 2KXQ, 1S7M, and 1LQL), the algorithm was halted prior to 1,000 generations. For PDBs 1EMN, 1NP6, and 1P9N, the algorithm completed the full 1,000 generations (parameters are listed in **Table S1**).

In each generation, the number of objective function evaluations was calculated by multiplying the population size (5 individuals) by the samples generated per individual (5) and the total number of islands (20), yielding 500 evaluations. In the 21^st^ island, we adjusted the evaluations to 25, resulting in 525 objective function calls per generation.

For proteins primarily composed of α-helices, GDT_TS distributions trended toward higher values, approaching 1, whereas RMSD_S distributions approached 0 (**Figure 1b, Table S2)**. This pattern indicates high structural similarity and stability among the designs, with minimal variance. However, in proteins 5UIU, 5YIY, and 2QQ8 (**Figure 1d-I, -J, -L)**, GDT_TS and RMSD_S distributions were more dispersed. This variance may stem from disordered regions at one terminus, which lack well-defined secondary structure and thus needed more generations for convergence.

Proteins dominated by α-helices showed narrow GDT_TS standard deviations (0.00 to 0.07) yet larger RMSD_S variation (0.04 to 0.42) (**Table S2**). Although higher RMSD_S variability helps prevent premature convergence, β-sheet proteins typically required longer runs and showed more structural diversity, evident in broader RMSD_S and GDT_TS distributions. Because β-sheet proteins had an average length of 32 residues – nearly double that of α-helical proteins (18 residues)-their search space as inherently more complex. Unstructured proteins posed even greater challenges for the algorithm.

We calculated the correlation between GDT_TS and RMSD_S for the top designs (**Figure 1c**). Data points are color-coded by secondary structure type: green (β-sheet), red (α-helix), and blue (unstructured). Red points cluster toward higher GDT_TS and lower RMSD_S values, whereas blue points show lower accuracy. Protein 2QQ8 (L), which contains an unstructured region, stands out with significantly lower GDT_TS and RMSD_S scores compared to other α-helical proteins.

Superimposing the highest-fitness designs onto their reference structures (**Figure 1d**) reveals notable structural similarity, particularly in α-helical and β-sheet regions. Proteins with unstructured segments showed lower similarity, although extended runs did improve alignment. Notably, the highest sequence identity between the best designs and the reference sequences was merely 24%, with an average of 11% (**Table S2**), suggesting that the algorithm can converge on structurally similar solutions despite low sequence identity.

On this same dataset of 23 proteins and by using a Bayesian ranking, we compared KCM with three well-known generative models: ProteinMPNN, ESM-IF1, and ProteinSolver. For a given target backbone, ProteinMPNN generated 256 sequences in four runs, while ESM-IF1 and Protein Solver each produced 250 sequences. For each target, we selected the sequence with the highest GDT_TS and the sequence with the lowest RMSD for comparison. For KCM, we likewise chose the best sequences according to each criterion. For a fair comparison, all solutions from each algorithm were sorted based on the fitness function used in our algorithm. When examining only 50 solutions (**Figures S1a-f**, and **Table S3**), KCM surpassed the other approaches in RMSD but lagged behind ESM-IF1 and ProteinMPNN in GDT_TS. When 250 solutions were considered (**Figures S1g-l**, and **Table S4**), KCM again outperformed all other methods in RMSD but fell short of ESM-IF1 in GDT_TS. Note that these proteins were used in the generative models’ training process. Deep learning algorithms typically perform better on data included in their training sets.

### Antimicrobial peptide design

As a proof-of-concept, we selected the small peptide IDR-2009 (sequence: KWRLLIRWRIQK), known for its potent antimicrobial activity^32^. We explored different combinations of the objective function components to determine whether they could yield soluble and synthesizable sequences.

We did not assume that the peptide’s function directly depends on its geometry. Instead, we investigated whether mirroring its predicted backbone, energy, and amino acid properties would yield new sequences with comparable structure and properties.

To design a new set of peptides using the IDR-2009 backbone as a template, we proceeded as follows: (i) We obtained the three-dimensional structure of the peptide using the Google Colab version of AlphaFold 2^37,38^, based on its sequence. Within AlphaFold 2, we used the default parameters but increased to 48 recycles and 3 relaxation iterations. This resulting backbone structure served as the input backbone for our KCM; (ii) We conducted the design under four different schemes, primarily varying the protein similarity function. The four schemes were:

#### All criteria included (AC)

All criteria in equation 2 were applied.

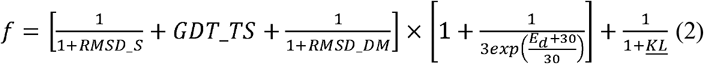

where *<u>KL</u>* is the mean Kullback–Leibler divergence (**equation 14**, see details in **Supplementary Information** section **S1.7. Similarity between descriptors**) of the descriptors computed from the sequence. RMSD_S, GDT_TS, and RMSD_CM are also defined in **Supplementary text**, in subsections **S1.1, S1.3**, and **S1.4**, respectively.

#### Descriptors not included (No Descriptors; ND)

We excluded the KL divergence terms from the objective function (equation 3).

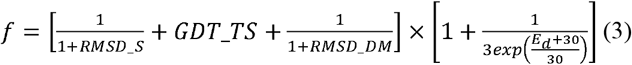

#### Mainly Descriptors (MD)

We excluded the geometric similarity criteria from the objective function and weighted the energy terms negligibly (equation 4).

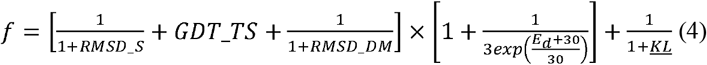

where *RMSD_S* = *RMSD_DM* = 10^6^ and *GDT_TS* = 0

#### No energy terms included (NE)

We omitted the energy term from the objective function (equation 5).

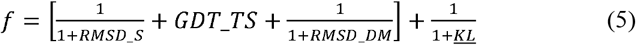

For each scheme, we generated a set of sequences and ranked them in descending order according to their fitness (**Table S5** lists the evolutionary algorithm parameters). We then analyzed these ranked sequences with AMPDiscover^39^ to predict their potential antimicrobial activity. We further examined the top 10 sequences from each scheme (**Tables S6–S8**).

### In vitro antimicrobial activity of peptides

We synthesized 12 sequences selected for their ease of synthesis and low aggregation potential: 5 from AC, 4 from NE, and 3 from MD. We excluded the ND group because its peptides were challenging to synthesize and solubilize. We evaluated their minimal inhibitory concentrations (MIC) against 11 clinically relevant strains, including the ESKAPEE pathogens. Nine of the 12 peptides (75%) exhibited MIC ≤64 μmol L^−1^ against at least one strain (**Figure 2a,b)**, surpassing hit rates from many existing ML/DL methods. The three inactive peptides (AC1, AC2, AC3) belonged to the AC group, had low net charge (<2), and displayed lower normalized hydrophobicity. This indicates that incorporating all criteria into the objective function might complicate the search landscape and hinder the discovery of highly active sequences. Further investigation is needed to confirm this hypothesis.

**Figure 2.**
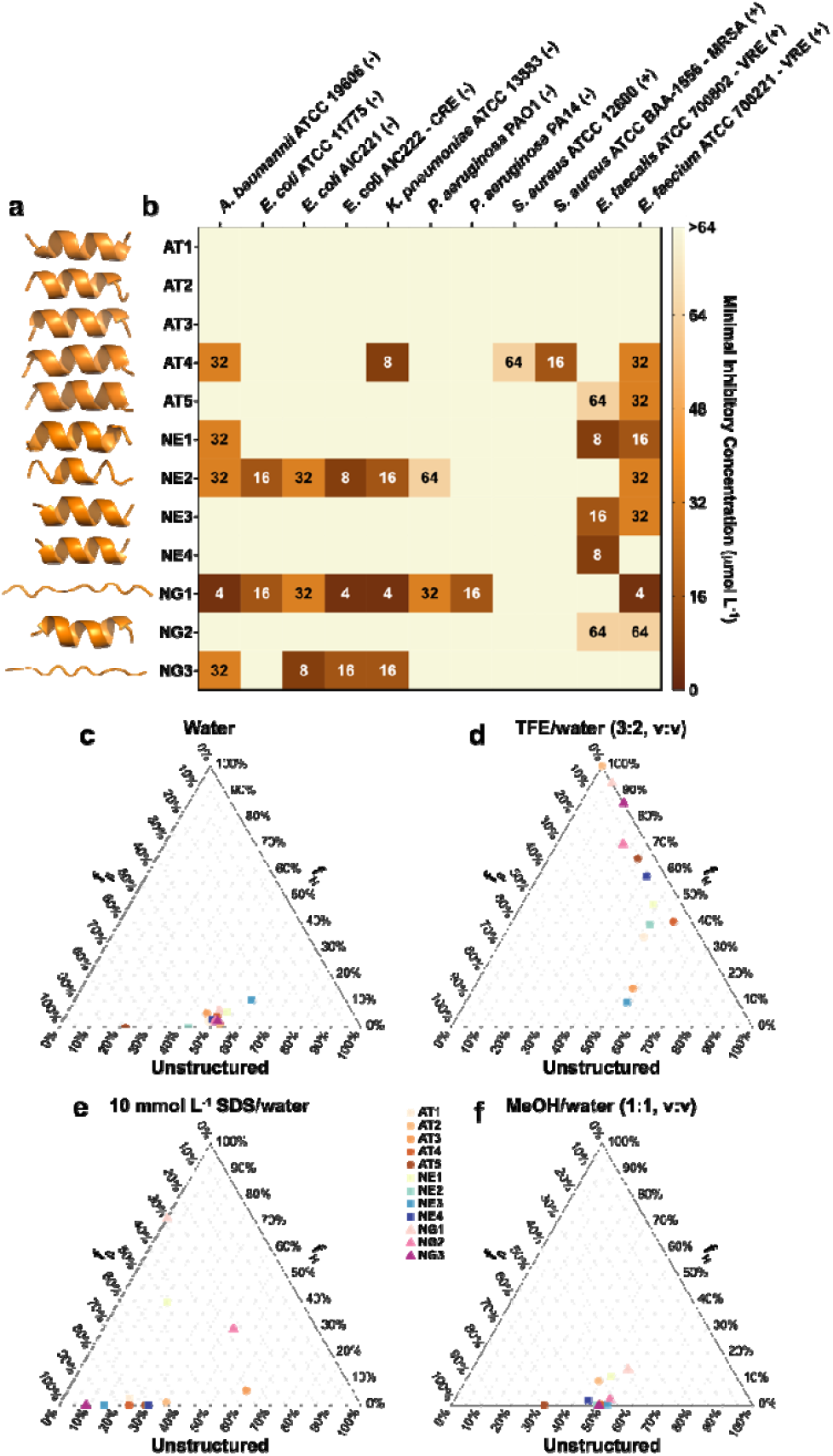
Antimicrobial activity and secondary structure of computationally designed peptides. **(a)** Backbone superposition of the reference protein (green) and the highest fitness designed peptides synthesized (orange) for experimental validation. **(b)** Heat map of the antimicrobial activities (μmol L^−1^) of the peptides against 11 clinically relevant pathogens, including four strains resistant to conventional antibiotics. Briefly, 10^6^ bacterial cells and serially diluted peptides (1-64 μmol L^−1^) were incubated at 37 °C. One day post-treatment, the optical density at 600 nm was measured in a microplate reader to evaluate bacterial growth in the presence of the peptides. MIC values in the heat map are the mode of the replicates in each condition. **(c-f)** Ternary plots showing the percentage of secondary structure for each peptide (at 50 μmol L^−1^) in four different solvents: **(c)** water, **(d)** 60% trifluoroethanol (TFE) in water, **(e)** Sodium dodecyl sulfate (SDS, 10 mmol L^−1^) in water, and **(f)** 50% methanol (MeOH) in water. Secondary structure fractions were calculated using the BeStSel server^48^.

### Secondary structure of designed peptides

Because the peptides were designed using different criteria, we sought to assess their secondary structure tendencies. To evaluate secondary structure, we subjected the peptides to four different media: water (**Figure 2c** and **Figure S2a**,**e**), a helix-inducing medium (trifluoroethanol in water, 3:2, v:v; **Figure 2d** and **Figure S2b,e**), a membrane-mimicking environment (sodium dodecyl sulfate, SDS, at 10 mmol L^−1^; **Figure 2e** and **Figure S2c,e**), as well as a β-inducer medium (methanol in water, 1:1, v:v; **Figure 2f** and **Figure S2d,e)**.

Interestingly, when geometric similarity criteria were omitted and energy terms were minimally weighted (MD group), the designed peptides displayed greater structural plasticity. They transitioned from unstructured forms in water and the methanol/water mixture to highly helical structures in the TFE/water mixture and predominantly β-like conformations in the membrane-mimicking medium. Peptides from the AC and NE groups remained unstructured in water yet adopted mostly β-like conformations in SDS and the methanol/water mixture. Notably, in the presence of SDS micelles, NE1 showed an equal proportion of β-like and helical structures.

Among the peptides, AC2, AC5, and NE4 formed the most helical conformations in the TFE/water mixture, despite being among the least amphiphilic peptides. Conversely, AC3 and NE3, which were among the more amphiphilic peptides, predominantly retained a β-like structure even in the helix-inducing medium. No clear trend emerged linking secondary structure directly to antimicrobial activity.

### Mechanism of action

We next explored the membrane-related mechanism of action of a subset of peptides (AC4, NE1, NE2, MD1, MD3) against *A. baumannii* and vancomycin-resistant *E. faecalis* (**Figure 3a**). Using NPN uptake assays (**Figure 3b** and **Figure S3a**) and membrane depolarization assays with DiSC_3_-5 (**Figure 3c,d** and **Figure S3b,c**), we found that all peptides disrupted the bacterial membrane potential to varying degrees. NE1 and MD3 were particularly effective at depolarizing the cytoplasmic membrane of *A. baumannii*. Although none outperformed polymyxin B in outer membrane permeabilization, several peptides matched levofloxacin’s effects, underscoring their potential as membrane-targeting agents.

**Figure 3.**
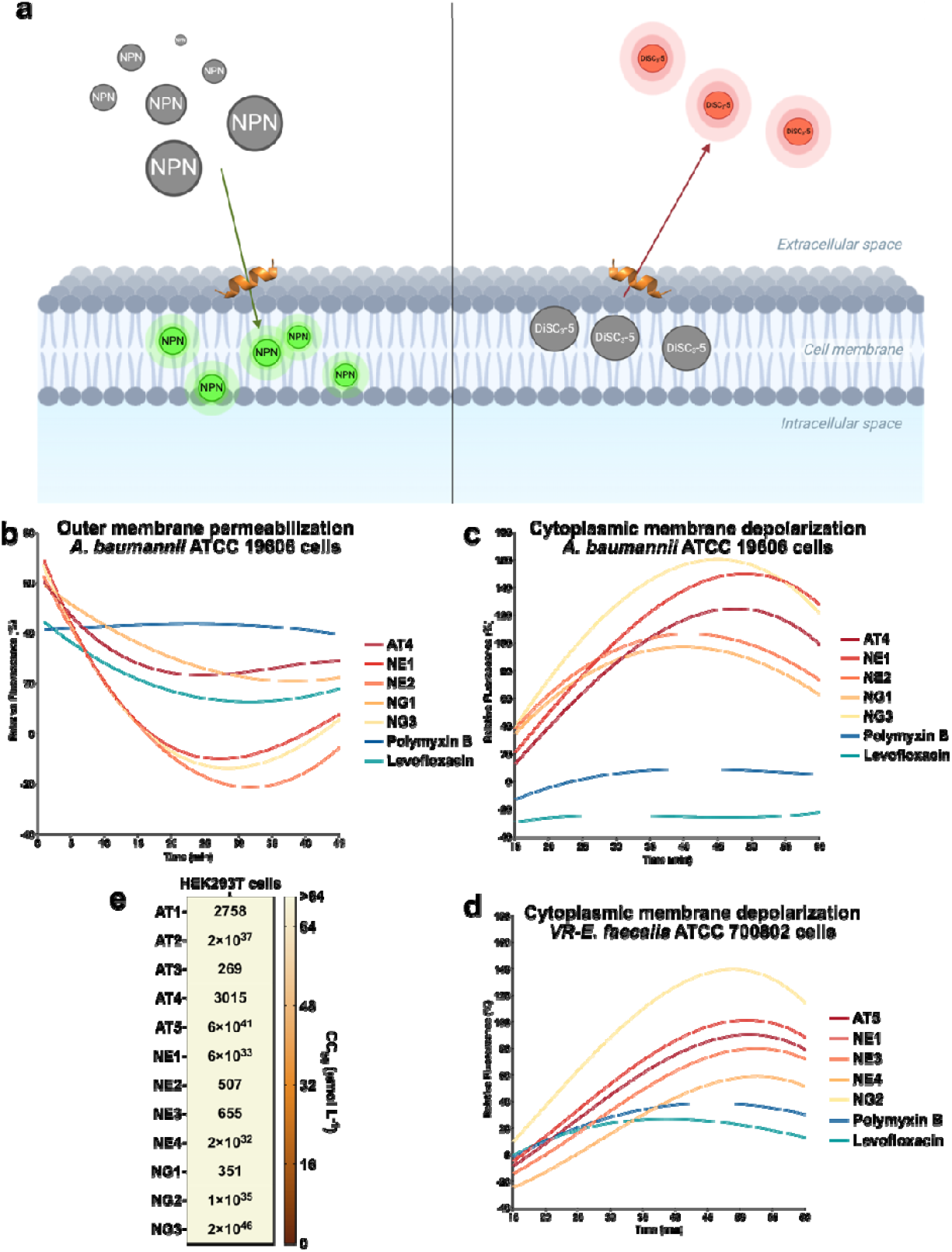
Mechanism of action and cytotoxic activity of designed peptides. To assess whether the peptides act on bacterial membranes, all active peptides against *A. baumannii* ATCC 19606 were subjected to outer membrane permeabilization and peptides active against *A. baumannii* ATCC 19606 and vancomycin-resistant *E. faecalis* ATCC 700802 were tested in cytoplasmic membrane depolarization assays. The fluorescent probe 1-(N-phenylamino)naphthalene (NPN) was used to assess membrane permeabilization **(a)** induced by the tested peptides. The fluorescent probe 3,3′-dipropylthiadicarbocyanine iodide (DiSC_3_-5) was used to evaluate membrane depolarization **(b)** caused by the designed peptides in *A. baumannii* ATCC 19606 and vancomycin-resistant *E. faecalis* ATCC 700802. The values displayed represent the relative fluorescence of both probes, with non-linear fitting compared to the baseline of the untreated control (buffer + bacteria + fluorescent dye) and benchmarked against the antibiotics polymyxin B and levofloxacin. **(c)** Cytotoxic concentrations leading to 50% cell lysis (CC_50_) were determined by interpolating the dose-response data using a non-linear regression curve. All experiments were performed in three independent replicates.

### Cytotoxicity assays

Cytotoxicity was evaluated using the human embryonic kidney (HEK293T) cells (**Figure 3e**)^40^. None of the 12 peptides caused significant cytotoxicity at the tested concentrations (4-64 μmol L^−1^), contrasting with the parental IDR-2009 peptide, which showed cytotoxicity^32^. The lowest CC_50_ among inactive peptides (AC3) was 269 μmol L^−1^, indicating a favorable therapeutic window.

### Anti-infective efficacy in animal models

We tested NE2 and MD1 in two murine models: skin abscess^41–44^ (**Figure 4a,b**) and a deep thigh infection^45–47^ (**Figure S4c–e**). In both models, the peptides reduced *A. baumannii* bacterial loads by up to two orders of magnitude, comparable to the last-resort antibiotic polymyxin B and to our other antibiotic control levofloxacin. No weight loss or skin damage was observed, indicating good tolerability. These results confirm that KCM-designed peptides can achieve clinically relevant anti-infective efficacy.

**Figure 4.**
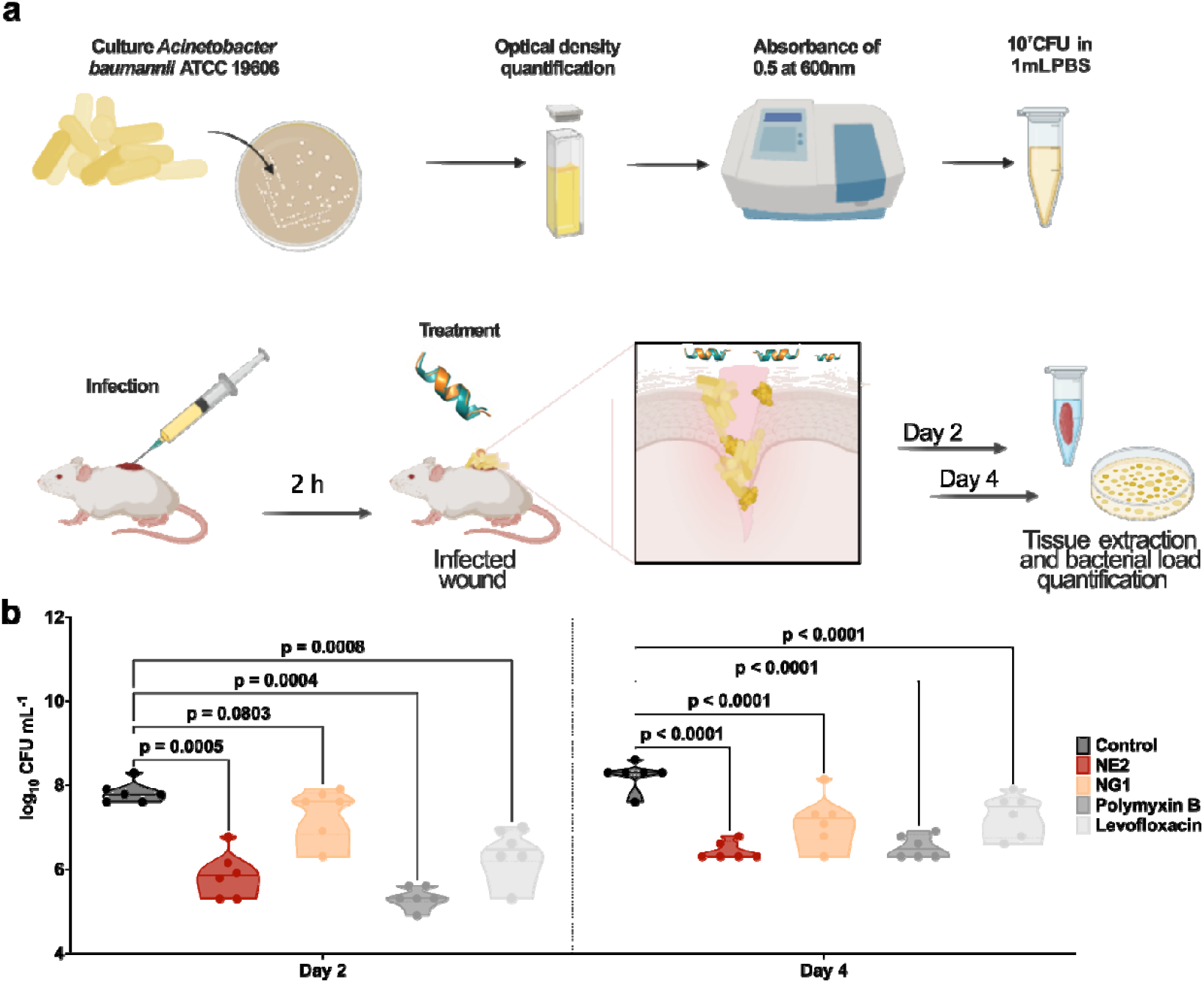
Anti-infective activity of designed peptides in a skin scarification animal model. **(a)** Schematic representation of the skin abscess mouse model used to assess the anti-infective activity of the designed peptides (n□=□6) against *A. baumannii* ATCC 19606. **(b)** NE2 and NG1, administered at their MIC (32 and 4 μmol L^−1^, respectively) in a single dose post-infection, inhibited the proliferation of the infection for up to 4□days after treatment compared to the untreated control group. Notably, NE2 reduced the infection in some mice, demonstrating activity comparable to the control antibiotics, polymyxin B and levofloxacin. Statistical significance in panel **b** was determined using one-way ANOVA followed by Dunnett’s test; P values are shown in the graphs. In the violin, the center line represents the mean, the box limits the first and third quartiles, and the whiskers (minima and maxima) represent 1.5□×□ the interquartile range. The solid line inside each box represents the mean value obtained for each group. Panel **a** was created with BioRender.com.

## Discussion

This study introduces and validates the KCM approach for computational protein and peptide design. In contrast to purely generative frameworks, which use trained models to rapidly generate large numbers of candidate sequences, KCM iteratively refines candidate sequences through structure prediction and optimization. Our results demonstrate that KCM can converge on sequences whose backbone geometries strongly resemble those of target proteins, spanning α-helices, β-sheets, and unstructured regions, even when sequence identity is low. This suggests that KCM effectively explores the structural landscape by continuously updating the distribution of candidate sequences based on both geometric and physicochemical criteria.

A key advantage of KCM is its ability to seamlessly integrate a wide range of objective functions beyond backbone geometry. In particular, it can incorporate Kullback–Leibler divergence or Jeffrey’s distance for amino acid descriptors, as well as energy terms and other chemically relevant properties. This flexibility allows researchers to add or remove objectives— such as solubility, stability, or functional motifs—within a single GPU environment, eliminating the need to retrain a deep learning model when new properties are introduced into the loss function. In contrast to generative models that require substantial computational resources for retraining, KCM offers an adaptable, resource-efficient solution for computational protein design.

Beyond benchmarking KCM on proteins of diverse secondary structure, we applied it to the design of novel antimicrobial peptides. Despite not explicitly encoding direct activity constraints into our fitness function, the KCM-derived peptides retained antimicrobial activity that was comparable to or exceeded many ML/DL-based designs reported in the literature. Several designs also exhibited lower cytotoxicity than the parental AMP (IDR-2009), demonstrating that KCM can discover sequences with improved therapeutic windows.

Structural analyses revealed that relaxing energy and geometric similarity constraints leads to higher structural plasticity, with designed sequences frequently transitioning between helical and β-like conformations depending on environmental factors. Notably, this plasticity did not necessarily reduce antimicrobial activity, suggesting that multiple structural pathways can yield effective membrane-disrupting peptides. The diversity of outcomes highlights the importance of balancing structural and energetic constraints with the capacity to explore new, potentially functional sequence space.

In summary, KCM provides an alternative paradigm for peptide and protein design, combining the adaptability of an iterative, optimization-based framework with the power of modern structure prediction tools. Its demonstrated success in designing peptides with potent antimicrobial activity and reduced toxicity highlights the method’s potential in therapeutic development. Future work may explore more sophisticated descriptors—such as immunogenicity, proteolytic stability, and post-translational modifications—as well as advanced multi-objective strategies to enhance both throughput and exploration of sequence space. Ultimately, the KCM model represents a promising step toward more versatile and customizable computational design methodologies for proteins and peptides.

### Limitations of the study

Despite these advantages, several challenges remain with the KCM model. Its reliance on iterative structure prediction makes it more computationally intensive than standard pre-trained generative approaches, such as ProteinMPNN or ESM-IF1, which can complete designs in seconds. For example, designing an α-helical sequence in our benchmark set requires about 12 hours on a single GPU (RTX 3090 Ti) coupled with an Intel i5 CPU with 32 GB of RAM. This task is particularly challenging for short, dynamic peptides as they may adopt multiple conformations in solution. Integrating ensemble-based structure prediction or incorporating biophysical data (e.g., nuclear magnetic resonance) could improve accuracy for more flexible targets.

Nonetheless, a key benefit of KCM is that it requires no training phase. If a new term is added to the objective function, there is no need to retrain any model, affording greater flexibility in exploring different design criteria. Improvements in parallelization or incorporating faster structure predictors could help address the remaining computational burden.

## Methods

### Objective function

The objective function that we propose in this work for the key-cutting machine approach consists of a combination of different similarity measures between the structure, energy and chemical/evolutionary descriptors of a designed protein and those of the target protein. For measuring structural similarity, we used the standard root mean square deviation (RMSD_S)^49,50^, the Global Distance Test Total Score (GDT_TS)^36^, the Template Modeling Score (TM_S)^51^ and the root mean square deviation between the corresponding distance matrices (RMSD_DM). The Rosetta energy function REF15^52^ is used to measure the energy of a protein structure. Lastly, the chemical descriptors computed consisted of a subset of the iLearn protein descriptors (**Table S9**)^11^ and the decoded vector representation from the ESM-2 language model^53^. Given the descriptor vectors for a model’s sequence and the reference’s sequence, the similarity between them can be computed by using Jeffrey’s distance^54^ (described in detail in **Supplementary Information**, section **S1. Similarity metrics**).

Notice that all these criteria measure the similarity or difference between a particular aspect of a designed protein and the reference protein. For example, RMSD_S, TM_S, GDT_TS, and RMSD_DM all measure structural similarity. In this case, the structure of a designed sequence is predicted using ESMFold, and the resulting structure is then superimposed with the reference structure. On the other hand, the Jeffrey’s distance between ESM-2 and iLearn descriptor vectors measures how similar the two proteins are in terms of both iLearn and ESM-2 descriptors.

The objective function to be maximized by our protein design algorithm is given as:

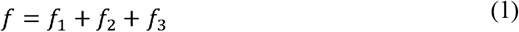

where,

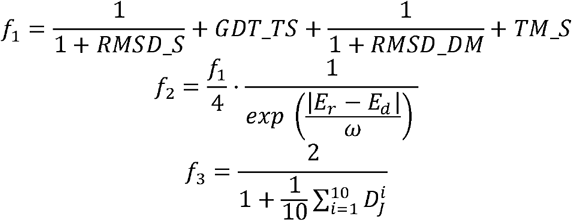

and *E*_*d*_ and *E* are, respectively of the model and reference structures, both calculated using the Rosetta energy function. Also, for all 1 ≤ *i* ≤ 10, the symbol 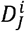 denotes the Jeffrey’s distance, computed from the th descriptor vector of both the model and the reference sequences. Note that the number of such vectors for each protein is 10, where 9 are computed with iLearn and 1 with ESM-2. Lastly, *ω ∈ R* is a parameter for contracting or dilating the exponential function.

The objective function is unaffected by rotations and translations of the structure, since both the model and reference structures are superimposed before calculating all criteria. Also, each of the terms *f*_1_, *f*_2_, and *f*_3_ contribute in a different magnitude to the total value of *f*. Since the maximum value given by said function is 7, then *f*_1_ represents 4/7 of the total value, while *f*_2_ and *f*_3_ represent 1/7 and 2/7 of the total, respectively. The function, will also have values close to zero when the model and reference proteins are different in all the criteria considered.

### Key-cutting Machine Algorithm

The algorithm follows the island paradigm^55^, and in total, there are 21 islands. The islands evolve synchronously and within each island we have a general procedure (**Figure 1**). The first component is a stochastic sequence generator (structure of the individuals in the population), which depends on the mathematical model designed to learn how highly fit sequences are distributed within the feasible solution space. The second component is a 3D structure predictor, in this case, ESMFold was used. The third component is the objective function, which is responsible for determining the similarity between the designs and the reference protein (the key). Finally, the fourth module is a repository where the top *t* sequences generated within the islands are stored according to their fitness.

The island model is equipped with a communication strategy. At the end of each generation, with probability p, two islands can collaborate, and the information shared is through the sequences. Each island stores the top t sequences with the highest fitness obtained throughout all generations of the algorithm. With probability proportional to their fitness, *m*_1_ sequences are selected from the t stored in the first island. All selected sequences that surpass the fitness of the weakest sequences in the second island will replace them. The same procedure is performed in reverse order.

Each island from 1 to 20 is equipped with a mechanism that operates as an EDA-type evolutionary algorithm (**Figure 1**). Additionally, the islands are disaggregated according to the structure of the individuals in the population, which will henceforth be referred to as the stochastic sequence generator. Each island from 1 to 20 will have *pop* individuals (stochastic generators) with the same structure. In turn, these generators depend on how the parameters that define them are updated. Furthermore, an additional island is introduced, where a traditional genetic algorithm is implemented (island #21).

The individuals on the islands are constructed from six stochastic models (**Table S10**). In Model 1, for each position in the sequence, the probability of occurrence of each amino acid is determined. The individuals on islands 1, 2, 3, and 4 are constructed based on this model. Models 2 to 4 include knowledge of amino acids, such as how they are grouped based on properties like polarity, propensity to form part of a specific secondary structure, or evolutionary similarity. In this way, three hierarchical models are constructed, where for each position in the sequence, the probability of belonging to one of the groups is determined first, and then, given the current group, the probability of belonging to the next group is determined, and so on until reaching the amino acid level. The individuals on islands 5 through 16 are constructed based on these models. Model 5 is a Bayesian network where the nodes represent the positions in the sequence. In this network, an arc is drawn from each position to the next position, reflecting the dependency between successive positions in the sequence. The individuals on islands 17 and 18 are constructed based on Model 5. Finally, Model 6 represents a Markov chain between each pair of consecutive positions in the amino acid sequence. In this model, the states correspond to the 20 amino acids and the individuals on islands 19 and 20 are constructed based on this model. Additionally, the update of the individuals’ parameters is carried out under 4 different schemes. In the first scheme (A, **Table S10**), the parameters (probabilities) are calculated based on the absolute frequencies of appearance obtained throughout the algorithm (islands 4, 8, 12, 16, 17, and 20). In the second scheme (B, **Table S10**), the parameters (probabilities) are calculated from the absolute frequencies of appearance but resetting these every *ngr* generations (islands 3, 7, 11, 15, 18, and 19). In the third scheme (C, **Table S10**), the parameters are calculated based on the probability proportional to the size of the accumulated value of the function *d*_*r*∗_ throughout the algorithm (islands 2, 6, 10, and 14). In the fourth scheme (D, **Table S10**), the parameters are calculated from the probability proportional to the size of the accumulated value of the function *d*_*r*∗_ but resetting this every *ngr* generations (islands 1, 5, 9, and 13). Where 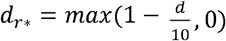 and *d* is the distance between the *C*_*α*_ atoms located at position *r*_α_ in the reference protein and the designed protein after superposition.

To update the parameters of the stochastic generator in generation *i*, two sets of sequences are selected. The first set contains *m* ∗ sequences, while the second set contains *t* ∗ sequences. The first set is formed from the *m* ∗ sequences with the highest fitness among the *m* sequences generated during generation *i* (*m* ∗ ≤ *m*). To construct the second set, *t* ∗ sequences are selected from the repository that contains the *t* sequences with the highest fitness generated in each island (*t* ∗ ≤ *t*). The selection of the *t* ∗ sequences is performed with a probability proportional to their fitness size (the details for the update rules of the stochastic generators are given in **Table S11** and **Supplementary Information**, section **S2. Stochastic generator of sequences**).

### Proteins to design

For testing the quality of the solutions produced by the KCM, we selected a small subset of proteins from the CATH Protein Structure Classification database, version 4.3.0, which is a free and publicly available online resource that provides information on the evolutionary relationships of protein domains^56^ and classifies them according to the secondary structures that compose them. This database is available at: http://download.cathdb.info/cath/releases/all-releases/v4_3_0/cath-classification-data/cath-domain-description-file-v4_3_0.txt.

In particular, we selected protein segments containing mainly ***α***-helices and ***β***-sheets. We also selected some proteins (**Table S12**) that did not have well defined secondary structures. In total, 23 proteins were selected, each between 15 to 35 residues long.

### Peptide Synthesis

All peptides for the experiments were obtained from pepMic and synthesized using solid-phase peptide synthesis with the Fmoc strategy. Peptides were purified using reverse-phase liquid chromatography (purity >95%).

### Bacterial strains and growth conditions

In this study, we used the following pathogenic bacterial strains: *Acinetobacter baumannii* ATCC 19606, *Escherichia coli* AIC221 [*Escherichia coli* MG1655 phnE_2::FRT (control strain for AIC 222)] and *Escherichia coli* AIC222 [*Escherichia coli* MG1655 pmrA53 phnE_2::FRT (polymyxin resistant; colistin-resistant strain)], *Klebsiella pneumoniae* ATCC 13883, *Pseudomonas aeruginosa* PAO1, *Pseudomonas aeruginosa* PA14, and *Staphylococcus aureus* ATCC 12600. All pathogens were grown in Luria-Bertani (LB) broth and on LB agar. Pseudomonas Isolation (*Pseudomonas aeruginosa* strains) agar plates were exclusively used in the case of *Pseudomonas* species. In all the experiments, bacteria were inoculated from one-isolated colony and grown overnight (16 h) in liquid medium at 37 °C. In the following day, inoculums were diluted 1:100 in fresh media and incubated at 37 °C to mid-logarithmic phase.

### Minimal inhibitory concentration assays

Broth microdilution assays^57^ were conducted to establish the minimum inhibitory concentration (MIC) for each peptide. Peptides were added to untreated polystyrene 96-well microtiter plates and serially diluted two-fold in sterile water, ranging from 1 to 64 μmol L□^1^. A bacterial inoculum at a concentration of 10 □ CFU mL□^1^ in LB medium was then mixed in a 1:1 ratio with the peptide solution. The MIC was determined as the lowest peptide concentration that completely inhibited bacterial growth after 24 h of incubation at 37 °C. Each assay was performed in three independent replicates.

### Circular dichroism experiments

The circular dichroism experiments were conducted using a J1500 circular dichroism spectropolarimeter (Jasco) in the Biological Chemistry Resource Center (BCRC) at the University of Pennsylvania. Experiments were performed at 25 °C, the spectra graphed are an average of three accumulations obtained with a quartz cuvette with an optical path length of 1.0 mm, ranging from 260 to 190 nm at a rate of 50 nm min^−1^ and a bandwidth of 0.5 nm. The concentration of all peptides tested was 50 μmol L^−1^, and the measurements were performed in water, a mixture of trifluoroethanol (TFE) and water in a 3:2 ratio, a mixture of methanol (MeOH) and water in a 1:1 ratiuo, and sodium dodecyl sulfate (SDS) in water at 10 mmol L^−1^, with respective baselines recorded prior to measurement. A Fourier transform filter was applied to minimize background effects. Secondary structure fraction values were calculated using the single spectra analysis tool on the server BeStSel^48^. Ternary plots were created in https://www.ternaryplot.com/ and subsequently edited.

### Outer membrane permeabilization assays

N-phenyl-1-napthylamine (NPN) uptake assay was used to evaluate the ability of the peptides to permeabilize the bacterial outer membrane. Inocula of *A. baumannii* ATCC 19606 were grown to an OD at 600 nm of 0.4 mL^−1^, centrifuged (10,000 rpm at 4 ºC for 10 min), washed and resuspended in 5 mmol L^−1^ HEPES buffer (pH 7.4) containing 5 mmol L^−1^ glucose. The bacterial solution was added to a white 96-well plate (100 μL per well) together with 4 μL of NPN at 0.5 mmol L^−1^. Consequently, peptides diluted in water were added to each well, and the fluorescence was measured at λ_ex_ = 350 nm and λ_em_ = 420 nm over time for 45 min. The relative fluorescence was calculated using the untreated control (buffer + bacteria + fluorescent dye) as baseline and the following equation was applied to reflect % of difference between the baselines and the sample:

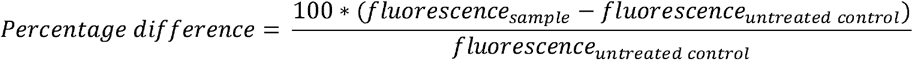

### Cytoplasmic membrane depolarization assays

The cytoplasmic membrane depolarization assay was performed using the membrane potential-sensitive dye 3,3’-dipropylthiadicarbocyanine iodide (DiSC_3_-5). *A. baumannii* ATCC 19606 in the mid-logarithmic phase were washed and resuspended at 0.05 OD mL^−1^ (optical value at 600 nm) in HEPES buffer (pH 7.2) containing 20 mmol L^−1^ glucose and 0.1 mol L^−1^ KCl. DiSC_3_-5 at 20 μmol L^−1^ was added to the bacterial suspension (100 μL per well) for 15 min to stabilize the fluorescence which indicates the incorporation of the dye into the bacterial membrane, and then the peptides were mixed 1:1 with the bacteria to a final concentration corresponding to their MIC values. Membrane depolarization was then followed by reading changes in the fluorescence (λ_ex_ = 622 nm, λ_em_ = 670 nm) over time for 60 min. The relative fluorescence was calculated using the untreated control (buffer + bacteria + fluorescent dye) as baseline and the following equation was applied to reflect % of difference between the baselines and the sample:

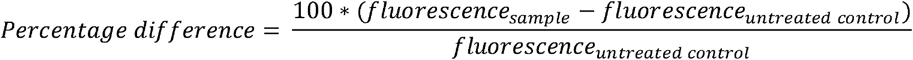

### Eukaryotic cell culture conditions

Human embryonic kidney (HEK293T) cells were obtained from the American Type Culture Collection (ATCC; CRL-3216™). The cells were cultured in high-glucose Dulbecco’s modified Eagle’s medium (DMEM) supplemented with 1% penicillin and streptomycin (antibiotics) and 10% fetal bovine serum (FBS) and grown at 37 °C in a humidified atmosphere containing 5% CO_2_.

### Cytotoxicity assays

One day before the experiment, an aliquot of 100 μL of the cells at 50,000 cells mL^−1^ was seeded into each well of the cell-treated 96-well plates used in the experiment (*i*.*e*., 5,000 cells per well)^40^. The attached HEK293T cells were then exposed to increasing concentrations of the peptides (4-64 μmol L^−1^) for 24 h. After the incubation period, we performed the (3-(4,5-dimethylthiazol-2-yl)-2,5-diphenyltetrazolium bromide) tetrazolium reduction assay (MTT assay)^9^. The MTT reagent was dissolved at 0.5 mg mL^−1^ in medium without phenol red and was used to replace cell culture supernatants containing the peptides (100 μL per well), and the samples were incubated for 4 h at 37 °C in a humidified atmosphere containing 5% CO_2_ yielding the insoluble formazan salt. The resulting salts were then resuspended in hydrochloric acid (0.04 mol L^−1^) in anhydrous isopropanol and quantified by spectrophotometric measurements of absorbance at 570 nm. All assays were done as three biological replicates.

### Skin abscess infection mouse model

The back of six-week-old female CD-1 mice under anesthesia were shaved and injured with a superficial linear skin abrasion made with a needle. An aliquot of *A. baumannii* ATCC 19606 (2×10^6^ CFU mL^−1^; 20 μL) previously grown in LB medium until 0.5 OD mL^−1^ (optical value at 600 nm) and then washed twice with sterile PBS (pH 7.4, 13,000 rpm for 3 min) was added to the scratched area. Peptides diluted in sterile water at MIC value were administered to the wound area 1 h after the infection. Two- and four-days post-infection animals were euthanized, and the scarified skin was excised, homogenized using a bead beater (25 Hz for 20 min), 10-fold serially diluted, and plated on McConkey agar plates for CFU quantification. The experiments were performed using three mice per group. The skin abscess infection mouse model was revised and approved by the University Laboratory Animal Resources (ULAR) from the University of Pennsylvania (Protocol 806763).

### Deep thigh infection mouse model

Six-week-old female CD-1 mice were rendered neutropenic by two doses of cyclophosphamide (150 mg Kg^−1^ and 100 mg Kg^−1^) applied intraperitoneally 3 and 1 days before the infection. At day 4 of the experiment, the mice were infected in their right thigh by a 100 μL intramuscular injection of the *A. baumannii* ATCC 19606 in PBS at concentration of 5×10^6^ CFU mL^−1^. The bacterial cells were grown in LB broth, washed twice with PBS solution, and diluted at the desired concentration. The peptides were administrated intraperitoneally two hours after the infection. Two-days post-infection mice were euthanized, and the tissue from the right thigh was excised, homogenized using a bead beater (25 Hz for 20 min), 10-fold serially diluted, and plated on McConkey agar plates for bacterial colonies counting. The experiments were performed using three mice per group. The deep thigh infection mouse model was revised and approved by the University Laboratory Animal Resources (ULAR) from the University of Pennsylvania (Protocol 807055).

## Acknowledgments

Cesar de la Fuente-Nunez holds a Presidential Professorship at the University of Pennsylvania and acknowledges funding from the Procter & Gamble Company, United Therapeutics, a BBRF Young Investigator Grant, the Nemirovsky Prize, Penn Health-Tech Accelerator Award, Defense Threat Reduction Agency grants HDTRA11810041 and HDTRA1-23-1-0001, and the Dean’s Innovation Fund from the Perelman School of Medicine at the University of Pennsylvania. Research reported in this publication was supported by the Langer Prize (AIChE Foundation), the NIH R35GM138201, and DTRA HDTRA1-21-1-0014. We thank Dr. Mark Goulian for kindly donating the following strains: *Escherichia coli* AIC221 [*Escherichia coli* MG1655 phnE_2::FRT (control strain for AIC222)] and *Escherichia coli* AIC222 [*Escherichia coli* MG1655 pmrA53 phnE_2::FRT (polymyxin resistant)]. We thank de la Fuente Lab members for insightful discussions. Figures created with BioRender.com are attributed as such. Molecules were rendered using the PyMOL Molecular Graphics System, Version 3.1.1 Schrödinger, LLC.

## Author contributions

Conceptualization: MDTT, YCL, CAB, CFN

Experimental methodology: MDTT

Computational methodology: YCL, CAO, CAB

Visualization: MDTT, YCL, CAB

Funding acquisition: CAB, CFN

Supervision: CAB, CFN

Software: YC, CAO, CAB

Formal analysis: MDTT, YCL, CAO, CAB

Writing – original draft: MDTT, YCL, CAB, CFN

Writing – review & editing: MDTT, YCL, CAB, CFN

Writing – review & editing final version of the paper: MDTT, YCL, CAB, CFN

## Data and materials availability

All training data, testing data, and code used to develop the machine learning model are freely available on GitHub (https://github.com/cbrizuel/KCM). All data pertaining to the experimental validation of generated peptides are available in the Supplementary Data or from the corresponding author upon reasonable request.

## Supplementary Information

### Supplementary text

#### S1. Similarity metrics

##### Notation

Let *A*_*m*_ and *A*_*r*_ denote respectively the set of atoms being considered in the model (designed) structure and the reference (or target) structure. Unless otherwise stated, we only consider the *Cα* atoms from each residue in the protein backbone. The number of atoms in a given structure is denoted as *n* ∈ *N*, where *n* = |*A*_*m*_| = |*A*_r_|. We will frequently use *a*_*m*,*i*_ ∈ *A*_*m*_ and *a*_*r*,*i*_ ∈ *A*_*r*_ to denote the *i* th pair of corresponding atoms in the model and reference structures. Furthermore, let *a*,*b* ∈ *A*_*m*_ ⋃ *A*_*r*_ be the Cartesian coordinates of two atoms, we denote the Euclidean distance between these atoms as *d*(*a*,*b*). We will also use *D*_*i*_ = *d*(*a*_*m*,*I*_, *a*_*r*,*i*_) as a shorthand.

##### S1.1. RMSD_S

The RMSD measures the average distance the atoms of the model structure deviate from their corresponding position in the reference structure^49,50^. The RMSD_S is measured in Angstroms and is computed after superimposing the two structures being compared. The RMSD_S is a positive real number, a value of zero indicates that the two structures are identical. The formula for calculating the RMSD_S is given by the following equation:

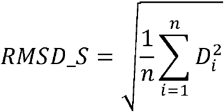

For calculating the RMSD_S value, we will consider not only the *C*_*α*_ atoms, but also the N, C and O atoms from each residue in the backbone.

##### S1.2. TM score

Like the RMSD, the TM Score also measures the average deviation distance between corresponding atoms in the model and reference structures after superimposing said structures. However, in the TM Score, the deviation distances are normalized by the number of atoms^51^. This allows the TM Score to describe more consistently the similarity between two structures regardless of their size. Also, since large deviation distances end up having a smaller contribution to the total score than short deviations, the TM Score is less sensitive than the RMSD to local misalignments in the structures. The TM Score is a real number in (0,1], where 1 indicates that the two structures are identical. The TM Score is given by the following formula:

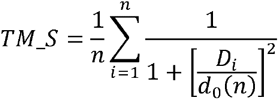

where *d*_0_(*n*) is the scaling factor determined by the size of the protein. This scaling factor is computed as:

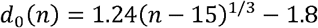

##### S1.3. GDT_TS

This score can be interpreted as the percentage of atoms in the model structure which deviate from their corresponding position in the reference structure in a distance that is within a given tolerance threshold (called cutoff distance)^36^. Usually, multiple cutoff distances are used, and the GDT_TS is the average of the percentages computed for each of these distances. The calculation of GDT_TS requires that the model and reference structures be superimposed. Being a percentage, the GDT_TS score is a real value in the interval [0,100], where 100 indicates that the two structures are identical under the smallest distance threshold.

Let, be the set of cutoff distances, and let *k* = |*S*|. Usually, the cutoff distances are *S* = {1,2,4,8}^36^. For all cutoff distances *s* ∈ *S* let *g*_s_: *N* → {0,1} be a function defined as follows:

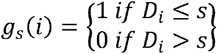

where 1 ≤ *i* ≤ *n*. Then, the GDT_TS is given by:

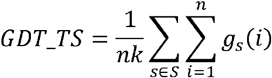

##### S1.4. RMSD_DM

The contact map of a protein refers to an alternative representation of the structure in which the distance between every pair of protein residues is described^58^. The contact map is a bidimensional symmetric matrix with binary values in which an entry value of 1 indicates that the distance between the corresponding residues is larger than a given contact threshold, and a value of zero indicates otherwise^59^. The same concept applies to the distance matrix, except that the Euclidean distances between each pair of atoms are directly stored as the matrix’s entries.

Let *A* be the set of atoms defining some protein structure. The distance matrix of *A*, denoted as *M*_*A*_ is the *n* × *n* matrix defined as:

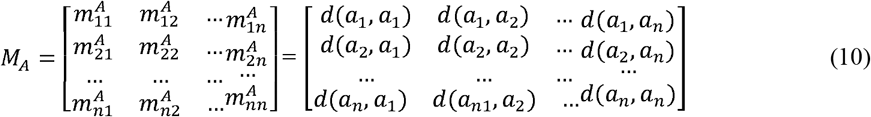

where, for all 1 ≤*i* ≤ *n*, the symbol *a*_*i*_ ∈ *A*, denotes the *i*th atom in the structure *A*.

The contact map can then be constructed from the distance map. Let *ε* ∈ *R* be the contact threshold, and let *h*_*ε*_: *R* → {0,1} be a function defined as:

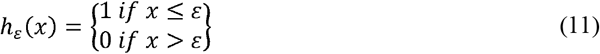

Then, the contact map of *A*, denoted as *C*_*A*_, is the *n* × matrix defined as:

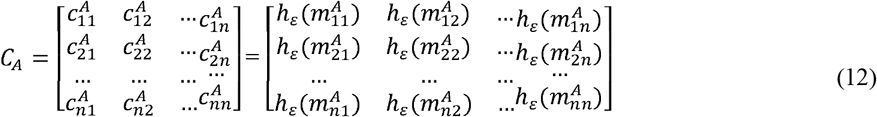

Given the distance maps from the model and reference structures, denoted 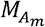 and 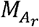 respectively, the RMSD_DM score is computed using the following formula:

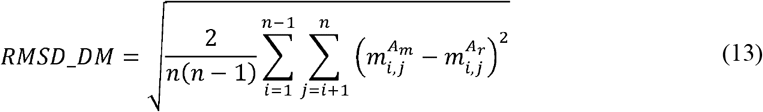

##### S1.5. ESM-2 descriptors

ESM-2 is a family of large Transformer-based neural networks that learned global protein properties and inter-residual relationships from known amino acid sequences^53,60^. These models were trained using a masking method, where random letters in the sequence were obscured and the neural network had to correctly predict them^61^. These models have proven to be useful for both computational protein design and protein structure prediction^62^. ESM-2 models are of particular interest in protein design because they can be used to produce proteins that do not belong to the collection of known proteins^63^.

The ESM-2 model family consists of six pre-trained models, each with 6, 12, 30, 33, 36, and 48 layers^53,64^. All models were trained with 65 million unique sequences. Given an amino acid sequence, any ESM-2 model returns as output a real-valued vector for each residue in the sequence. The protein and per-residue properties and the inter-residual relationships learned by the model is encoded in this vector of descriptors^53,64^. Depending on the number of layers used, each model produces larger or smaller descriptor vectors. For example, the 33-layer model produces descriptor vectors with 1,280 values per residue, while the 48-layer model produces 5,120 values per residue.

##### S1.6. iLearn descriptors

iLearn is a software toolkit that integrates several algorithms and procedures for calculating, extracting, analyzing and selecting protein and nucleotide features^65^. A subset of nine of these feature calculators (mainly those describing physicochemical characteristics of the amino acid sequence) was selected to be utilized in our KCM (**Table S9**).

##### S1.7. Similarity between descriptors

Given the descriptor vectors for a model’s sequence and the reference’s sequence, the similarity between them can be computed by using the Kullback-Leibler divergence and the Jeffrey’s distance^54^. Given two probability distributions, *P* and *Q* defined over the same set of data points *X*, the Kullback-Leibler divergence, denoted *D*_*KL*_(*P*||*Q*), measures how much distribution *P* diverges from the reference distribution *Q*. On the other hand, Jeffrey’s distance, denoted *D*_*J*_ is simply a symmetric version of the Kullback-Leibler divergence^54^. In the context of our protein design algorithm, distribution *P* would be obtained from the iLearn and ESM-2 descriptor vectors of a model sequence, and distribution *Q* would be obtained from the iLearn and ESM-2 descriptor vectors of the reference sequence.

Kullback-Leibler divergence is computed as^54^:

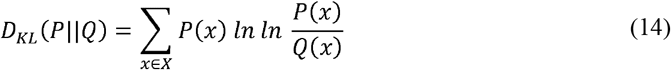

Meanwhile, Jeffrey’s distance is simply computed as :

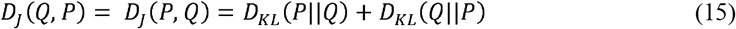

All the selected iLearn feature calculators produce real-valued descriptor vectors (**Table S9)**. Some of these vectors (like the ones produced by GDPC and GTPC) contain relative frequency values in [0,1], while others (like the ESM-2 descriptor vectors) contain absolute values, absolute frequency or percentage point values. In the first case, the relative frequency values are used directly as the probability distribution *P* or *Q* for calculating Jeffrey’s distance. In the second case, a histogram is computed, dividing the descriptor values’ range into 20 equal-sized subintervals and counting the number of values in each subinterval. These numbers are then converted into relative frequency values that are used as the distribution *P* or *Q* for calculating Jeffrey’s distance.

#### S2. Stochastic generator of sequences

The stochastic sequence generator is based on six mathematical models designed to learn how the highest fitness sequences are distributed within the feasible solution space. The generator also depends on the procedure used to update the parameters defined in the models. The first four are hierarchical models that learn the distribution of amino acids at each position in the sequence, without considering the relationships between different positions. To address these relationships, two models are employed: a Bayesian network and a Markov chain, which account for the interactions between different positions in the sequence.

The hierarchical models classify amino acids into different groups according to certain properties that they share with others. These properties are polarity, propensity to forming a certain secondary structure^66,67^, and structural and evolutionary similarity^65,68^. The latter refers to the fact that the substitution of one amino acid for another has a higher probability of being accepted if the two residues belong to the same group. The amino acids classification into different groups is as follows. The groups according to their polarity are:

- Hydrophilic: *C*_11_ = {*K, R, H, N, E, D, Q*}
- Hydrophobic: *C*_12_ = {*F, I, M, V, L*}
- Moderate polarity: *C*_13_ = {*Y, W, T, S, P, A, G, C*}

The groups according to the propensity of forming different secondary structures are:

- α-helices: *C*_21_ = {*L*,*A*,*Q*,*M*,*E*,*R*,*K*}
- β-sheet: *C*_22_ ={*S*,*N*,*D*,*P*,*G*}
- Loops: *C*_23_ = {*Y*,*T*,*W*,*F*,*I*,*V*}
- Others: *C*_24_={*H*,*C*}

The groups according to their evolutionary relationship are:

- *C*_31_ = {*Y*,*F*,*W*}
- *C*_32_ = {*R*,*K*,*H*}
- *C*_33_ = {*D*,*Q*,*N*,*E*}
- *C*_34_ = {*V*,*L*,*I*,*M*}
- *C*_35_ = {*S*,*G*,*P*,*T*,*A*}
- *C*_36_= {*C*}

Lastly, the groups according to their structural similarity are:

- *C*_41_= {*D*,*E*},*C*_42_ = {*N*,*Q*}, *C*_43_ = {*V, L*], *C*_44_ = {*S*,*T*},
- *C*_45_ - {*A, G*}, *C*_46_ = {*K*}, *C*_47_ = {*R*}, *C*_48_ = {*H*}, *C*_49_= {*I*}*C*_4,10_ = {*M*}, *C*_4,11_= {*F*}, C_4,12_ = {*Y*},
- *C*_4,13_ = {*W*}, *C*_4,14_ = {*P*}, *C*_4,15_ = {*C*}

Model 1 is the more general model and does not assume any previous knowledge about the amino acids or how they can be grouped together. Thus, for each position in the sequence, only the probability that a certain amino acid occupies that position is determined in this model. Models 2, 3, and 4 must determine the probability that a given amino acid belongs to one of such groups, and then, given a certain group, determine the probability of belonging to the next group. The order in which the different groups are visited is as follows:

- Model 2:
- Model 3:
- Model 4: where denotes each of the 20 standard amino acids, and ______ ______ ______ ______. Table S11 lists the parameters to be estimated by each model, 1 through 4.

In a Bayesian network, the number of parameters to be estimated increases with the number of arcs that are defined and, consequently, the number of samples needed to determine the parameters is higher. Thus, a Bayesian network is defined as follows: each position of the sequence corresponds to a node of the network, an arc is established between two nodes if they are in consecutive positions and the arc will point towards the node that is in a higher position. Model 5 will be a Bayesian network and the parameters it must estimate are listed in **Table S11**.

Matrix ***P***^***q***^can be thought of as the transition array at time 1 of a Markov chain. Also, given the nature of such transition matrix, the Markov chain would have a stationary distribution, and, consequently, the transition matrix can be determined when the time tends to infinity. In this way, is used to estimate the amino acids in the sequence. Model 6 will be a Markov chain that needs to estimate the parameters shown in **Table S11**.

The stochastic sequence generator depends on the method used to estimate and update the parameters in each generation. Therefore, for each parameter in models 1 to 4, with fixed *i, j* and *r*, we have the arrays (type A) or (type B), depending on the approach used. Both arrays initially have value 1 in their positions and, depending on the case, the probability assigned to each parameter is proportional to its corresponding value stored in the arrays and.

The arrays or are updated at the end of each generation, and the parameters are then recalculated. The process of updating the parameters will be explained with an example: assume that, on a given island, the model *m=4* is being used to learn how the amino acids are distributed at each position of the sequence. Also assume that, at position *r*_^*^_ of the sequence number selected to be updated, there is amino acid and and, where ______, ______, ______, and ______. Under these conditions, the arrays are updated as follows:

where 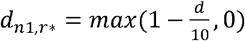 and *d* is the distance between the atoms of the corresponding amino acid in the reference sequence after both proteins are superimposed.

In this way, updating the arrays {*A*,*B*}^*ijr*^ is performed for the amino acids corresponding to all the positions of all the sequences selected for updating their parameters. The probabilities of each parameter are then calculated proportionally to the values stored in the arrays {*A*,*B*}^*ijr*^.

Additionally, the information stored in type A arrays corresponds to the frequency of appearance of amino acids or groups in the positions. Meanwhile, the information stored in type B is more selective, since, if the amino acid appears in that position but after superposition of the *C*_*α*_ atoms are distanced more than 10 Å away, then the information stored regarding that amino acid or group is negligible.

Due to the nature of models 5 and 6, type A arrays are always used to update their associated parameters. That is, the stored information corresponds to the absolute frequencies of the amino acids. In this way, all the tools are already given to explain how the islands are disaggregated according to the stochastic generator and the way the generator is used to update and calculate the parameters.

Four islands were built for each of the models 1 to 4. Two of these islands use type A arrays to determine the parameters, while the other two use type B arrays. Likewise, the two islands that use the same type of array differ in that the corresponding values in each array in one of the models are incremented throughout the algorithm, while in the other model, the values are reset every *ngr* generations. For models 5 and 6, only two islands were built, both of which use type A arrays. As in the previous case, these islands are different in that the values stored in those arrays are reset every *ngr* generations. In total, 20 islands were built that feature the stochastic sequence generator, while island 21 implements a traditional genetic algorithm explained above.

## Supplementary Tables

**Table S1.** Parameter values of KCM used in the design of selected proteins. *ng*: Maximum number of generations; *gdt*: GDT_TS threshold used as a stopping criterion; *s*: Number of islands; *p*: Information exchange rate between islands; *pop*: Population size, *t*: Number of elite sequences stored on each island; *m*□: Number of sequences selected for migration between islands; *m*: Number of sequences sampled at each generation; *ngr*: Number of generations after which parameters are reset for islands 1, 3, 5, 7, 9, 11, 13, 15, 17, 18, and 19; *m*^*^: Number of best sequences (from the m samples) used for parameter updates; *t*^*^: Number of best sequences (from the t stored sequences on each island) used for parameter updates.

| Parameter | Value |
| --- | --- |
| *ng* | Dependent of the protein |
| *gdt* | 0.9 for β-sheets and 1 for the rest |
| *p* | 0.1 |
| *pop* | 5 |
| *t* | 150 |
| *s* | All 21 islands |
| *m* | 5 |
| *ml* | 1 |
| *ngr* | 10 |
| *m\** | 3 |
| *t\** | 2 |

**Table S2.** Statistics of the 100 best designs for each protein. 2D: is the corresponding secondary structure, AH: alpha helix, BS: beta sheets, ND: not defined, L: character corresponding to each protein in Figure 3, PDB id.: the four letter protein ID in the PDB, RMSD_S Min, Mean and Sd: minimum, mean value and standard deviation for RMSD_S of the 100 designs of best fitness for each protein, GDT_TS Max, Mean and Sd: maximum, mean value and standard deviation for GDT_TS corresponding to the 100 best fitness for each protein, Identity: is the identity percentage of the highest fitness design with the reference sequence, n: length of the selected subsequence from the protein, and *ng*: number of generations employed in the design.

| 2D | L | RMSD_S |  |  | GDT_TS |  |  | Identity (%) | N | ng |
| --- | --- | --- | --- | --- | --- | --- | --- | --- | --- | --- |
|  |  | Min | Mean | Sd | Max | Mean | Sd |  |  |  |
| BS | A | 1.45 | 2.01 | 0.21 | 0.79 | 0.73 | 0.03 | 11 | 27 | 1,000 |
| BS | B | 1.19 | 2.05 | 0.40 | 0.91 | 0.73 | 0.07 | 12 | 26 | 210 |
| BS | C | 2.05 | 2.52 | 0.22 | 0.75 | 0.68 | 0.03 | 4 | 27 | 1,000 |
| BS | D | 1.10 | 2.00 | 0.38 | 0.92 | 0.74 | 0.05 | 12 | 26 | 305 |
| BS | E | 1.02 | 1.89 | 0.19 | 0.91 | 0.74 | 0.04 | 17 | 23 | 250 |
| BS | F | 1.61 | 2.17 | 0.23 | 0.84 | 0.70 | 0.04 | 10 | 30 | 1,000 |
| BS | G | 1.11 | 1.49 | 0.14 | 0.90 | 0.82 | 0.03 | 7 | 28 | 890 |
| BS | H | 2.27 | 2.64 | 0.22 | 0.72 | 0.66 | 0.03 | 3 | 33 | 1,000 |
| AH | I | 0.80 | 1.56 | 0.36 | 0.97 | 0.86 | 0.06 | 11 | 19 | 100 |
| AH | J | 0.61 | 1.67 | 0.42 | 0.98 | 0.84 | 0.07 | 11 | 19 | 100 |
| AH | K | 0.40 | 0.66 | 0.11 | 1.00 | 0.98 | 0.02 | 20 | 20 | 100 |
| AH | L | 2.19 | 2.82 | 0.29 | 0.70 | 0.59 | 0.04 | 6 | 19 | 100 |
| AH | M | 0.28 | 0.41 | 0.04 | 1.00 | 0.99 | 0.00 | 5 | 17 | 100 |
| AH | N | 0.47 | 0.74 | 0.13 | 0.99 | 0.97 | 0.01 | 20 | 20 | 100 |
| AH | O | 0.48 | 0.58 | 0.06 | 0.99 | 0.99 | 0.00 | 12 | 17 | 100 |
| AH | P | 0.47 | 0.61 | 0.07 | 1.00 | 0.98 | 0.01 | 0 | 19 | 100 |
| AH | Q | 0.41 | 0.69 | 0.13 | 1.00 | 0.97 | 0.02 | 16 | 19 | 100 |
| AH | R | 0.30 | 0.38 | 0.05 | 1.00 | 0.99 | 0.00 | 0 | 17 | 100 |
| AH | S | 0.54 | 0.85 | 0.19 | 0.99 | 0.95 | 0.02 | 24 | 17 | 100 |
| AH | T | 0.88 | 1.19 | 0.14 | 0.95 | 0.89 | 0.03 | 17 | 18 | 100 |
| ND | U | 2.01 | 2.58 | 0.22 | 0.70 | 0.61 | 0.04 | 13 | 15 | 100 |
| ND | V | 2.01 | 2.83 | 0.33 | 0.70 | 0.59 | 0.05 | 0 | 15 | 100 |
| ND | W | 2.09 | 2.87 | 0.34 | 0.72 | 0.59 | 0.05 | 20 | 15 | 100 |

**Table S3.** Probability of belonging to each region, resulting from the sign test (Left, Upper, and Right). Best 50 solutions. Left is the probability of KCM losing against the other methods, Upper is the probability of equal performance, and Right is the probability of KCM wining.

| Region | ESM-IF1 |  | Protein Solver |  | ProteinMPNN |  |
| --- | --- | --- | --- | --- | --- | --- |
|  | GDT_TS | RMSD_S | GDT_TS | RMSD_S | GDT_TS | RMSD_S |
| Left | 0.3158 | 0.49245 | 0.00 | 0.00 | 0.17835 | 0.404 |
| Upper | 0.4892 | 0.0126 | 0.0045 | 0.0001 | 0.64405 | 0.00315 |
| Right | 0.195 | 0.49495 | 0.9955 | 0.9999 | 0.1776 | 0.59285 |

**Table S4.** Probability of belonging to each region, resulting from the sign test (Left, Upper, and Right). Best 250 solutions. Left is the probability of KCM losing against the other methods, Upper is the probability of equal performance, and Right is the probability of KCM wining.

| Region | ESM-IF1 |  | Protein Solver |  | ProteinMPNN |  |
| --- | --- | --- | --- | --- | --- | --- |
|  | GDT_TS | RMSD_S | GDT_TS | RMSD_S | GDT_TS | RMSD_S |
| Left | 0.3183 | 0.3176 | 0 | 0 | 0.08085 | 0.3203 |
| Upper | 0.48485 | 0.01245 | 0.0051 | 0 | 0.7694 | 0.0131 |
| Right | 0.19685 | 0.66995 | 0.9949 | 1 | 0.14975 | 0.6666 |

**Table S5.** Parameter values used in KCM to design IDR 2009.

| Parameter | Value |
| --- | --- |
| ng | (A)-1770, (B)-1160, (C)-1080, (D)-470 |
| gdt | 1 |
| p | 0.10 |
| pop | 5 |
| t | 80 |
| s | all the islands except island 21 |
| m | 3 |
| m1 | 1 |
| ngr | 8 |
| m* | 2 |
| t* | 1 |

**Table S6.**
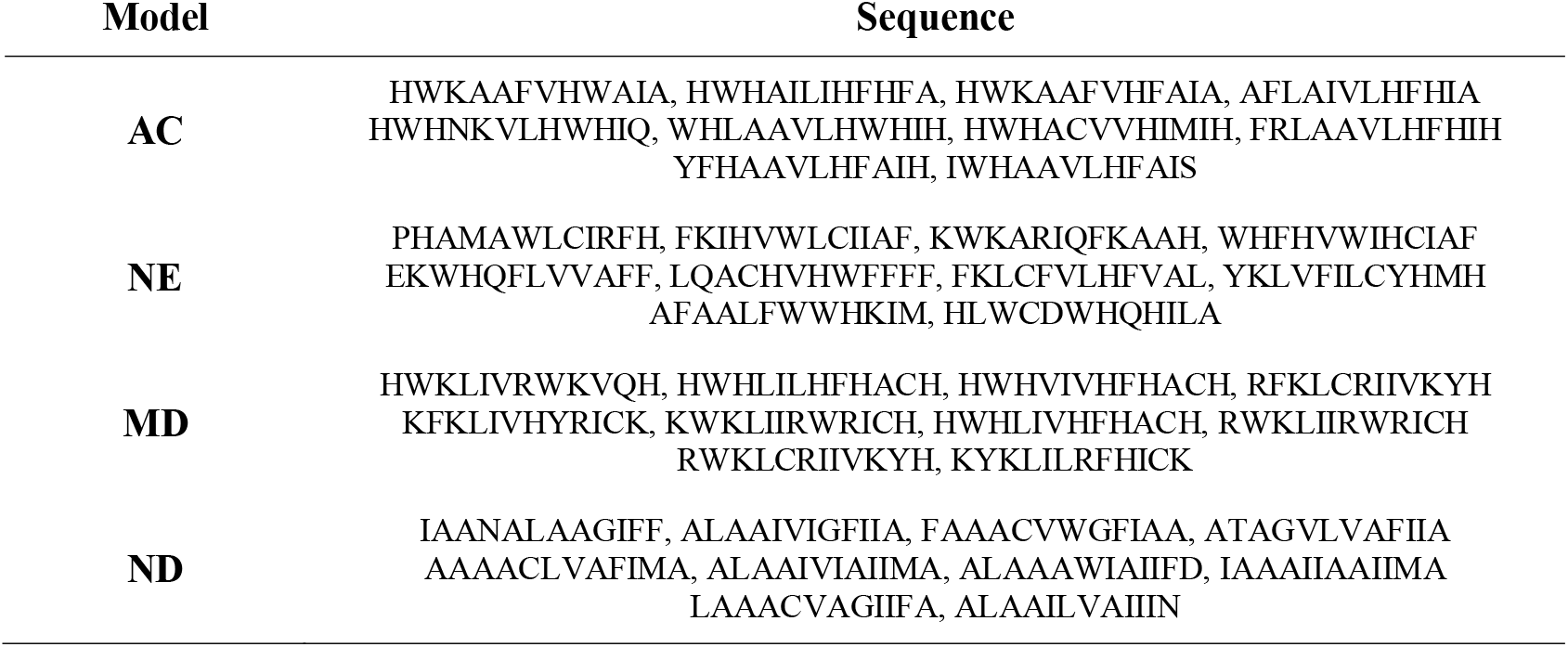
Ten best designs per combination of objective function terms (target IDR-2009).

**Table S7.**
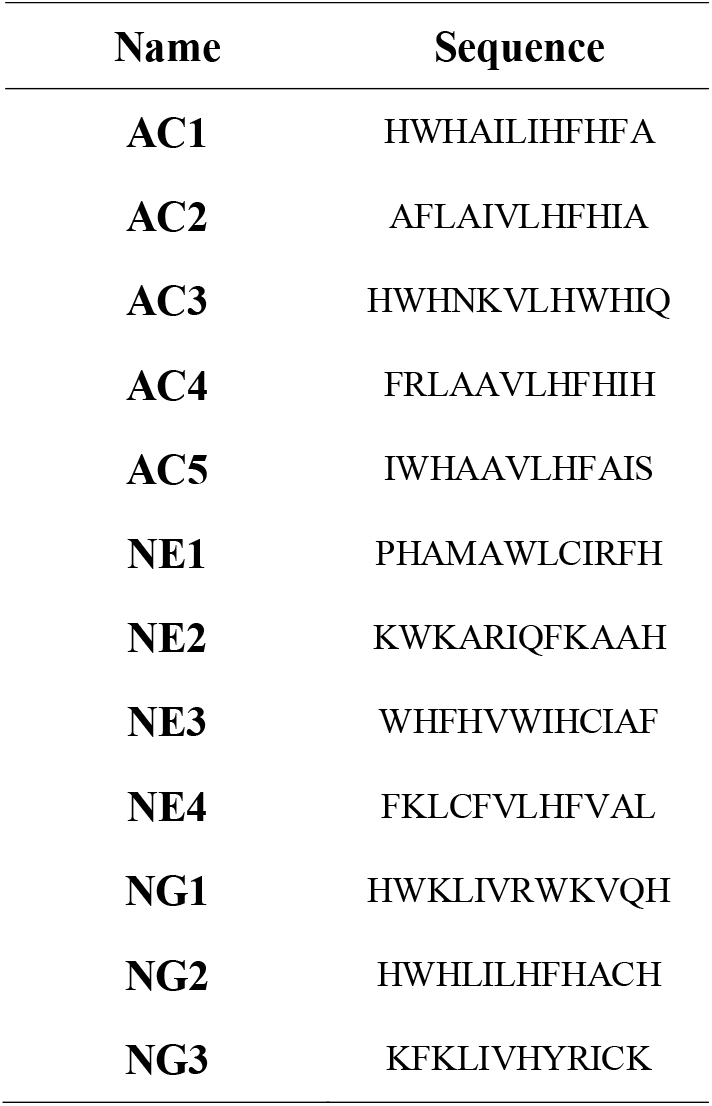
Selected peptides for experimental validation.

**Table S8.** Similarity criteria for the synthesized peptides using the backbone of IDR 2009 as a template.

| **ID** | **Energy** | **RMSD_S** | **GDT_TS** | **Descriptors similarity** | **Identity (%)** |
| --- | --- | --- | --- | --- | --- |
| AC1 | 29.68 | 0.36 | 1.00 | 0.89 | 8 |
| AC2 | 24.07 | 0.27 | 1.00 | 0.84 | 0 |
| AC3 | 120.88 | 0.30 | 1.00 | 0.89 | 8 |
| AC4 | 33.75 | 0.29 | 0.99 | 0.87 | 0 |
| AC5 | 19.85 | 0.29 | 0.99 | 0.82 | 8 |
| NE1 | 55.04 | 0.28 | 0.99 | 0.83 | 0 |
| NE2 | 79.29 | 0.69 | 0.98 | 0.87 | 25 |
| NE3 | 43.50 | 0.39 | 1.00 | 0.83 | 8 |
| NE4 | 61.95 | 0.35 | 1.00 | 0.81 | 0 |
| MD1 | 44.00 | 5.54 | 0.34 | 0.98 | 42 |
| MD2 | 38.07 | 0.65 | 0.97 | 0.97 | 17 |
| MD3 | 59.48 | 5.52 | 0.33 | 0.97 | 42 |
| IDR 2009 | 35.12 | 0.00 | 1.00 | 1.00 | 100 |

**Table S9.**
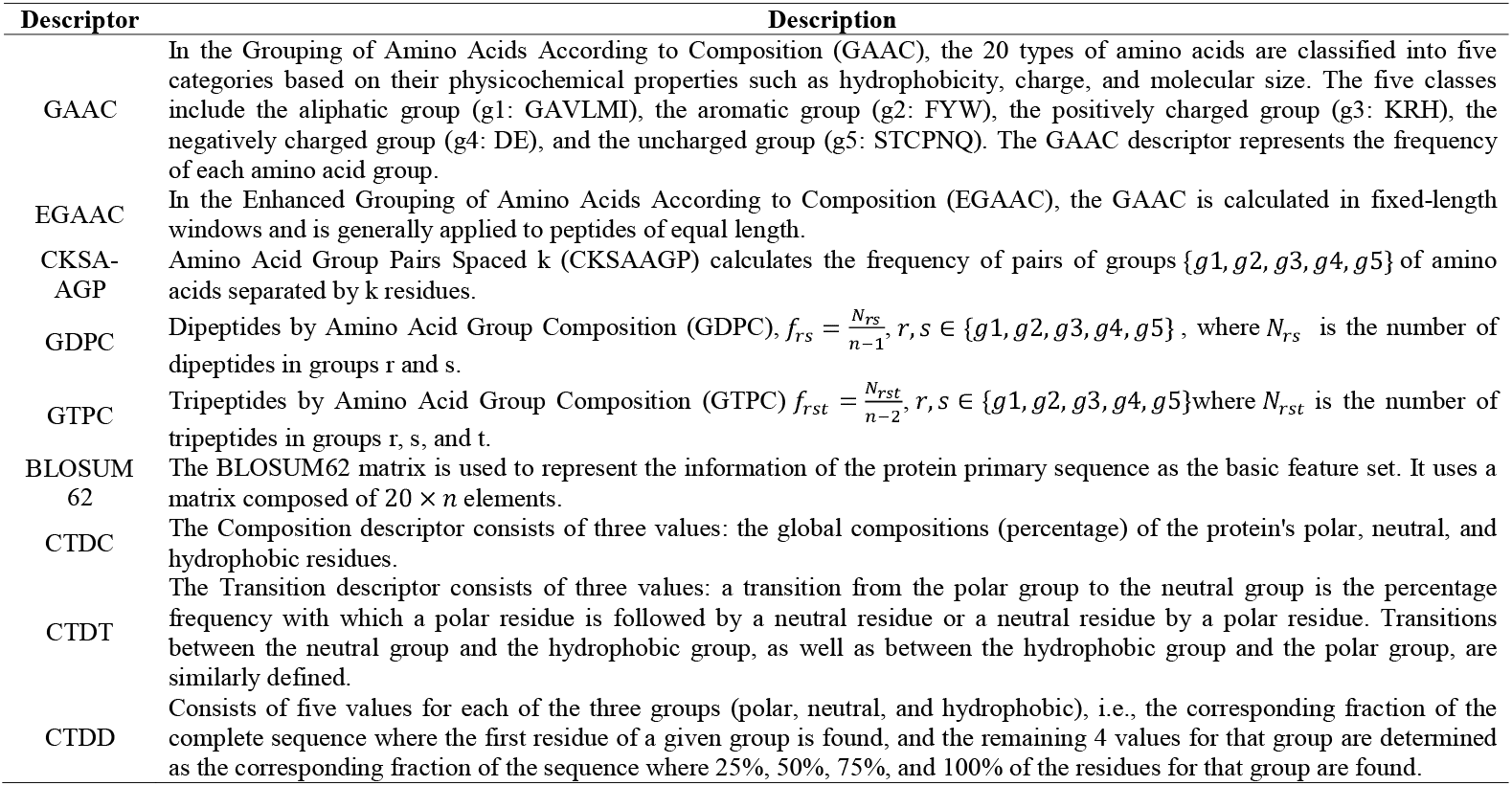
Description of the selected iLearn descriptors.

**Table S10.** Combination of models and parameters update strategies used in every island. (A) Absolute frequencies of appearance across all generations. (B) Absolute frequency of appearance reset every ngr generations. (C) Cumulative value of the function *d*_*r*∗_ across all generations. (D) Cumulative value of the function *d*_*r*∗_ reset every *ngr* generations. Where 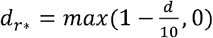 and *d* is the distance between the *C*_*α*_ atoms located at position *r*∗ in the reference protein and the designed protein after superposition.

| Parameter update strategy | Model 1 | Model 2 | Model 3 | Model 4 | Model 5 | Model 6 |
| --- | --- | --- | --- | --- | --- | --- |
| A | 4 | 8 | 12 | 16 | 17 | 20 |
| B | 3 | 7 | 11 | 15 | 18 | 19 |
| C | 2 | 6 | 10 | 14 | - | - |
| D | 1 | 5 | 9 | 13 | - | - |

**Table S11.**
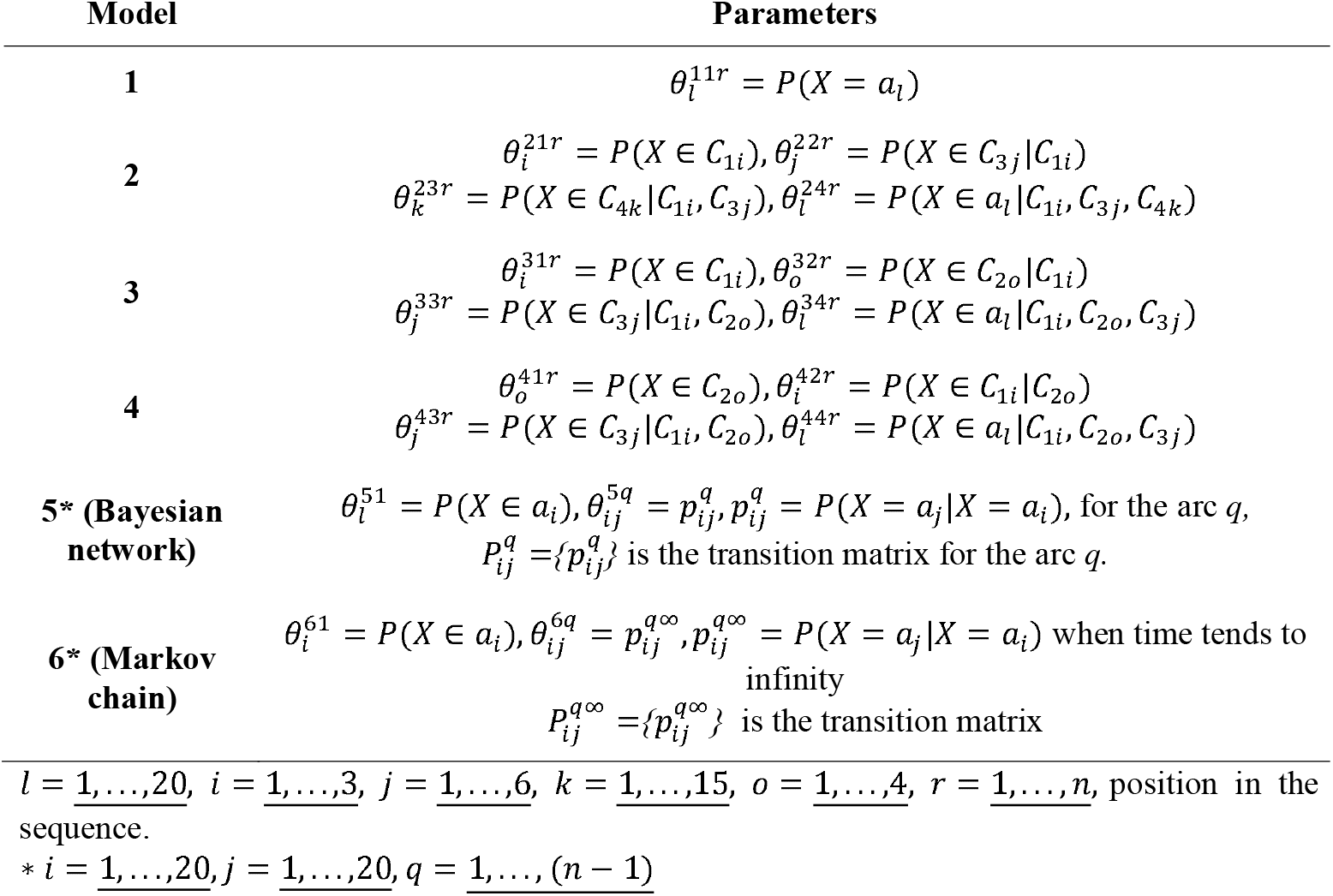
The parameters associated with models, 1 through 4 in each sequence position, and with Bayesian network and Markov chain models.

**Table S12.**
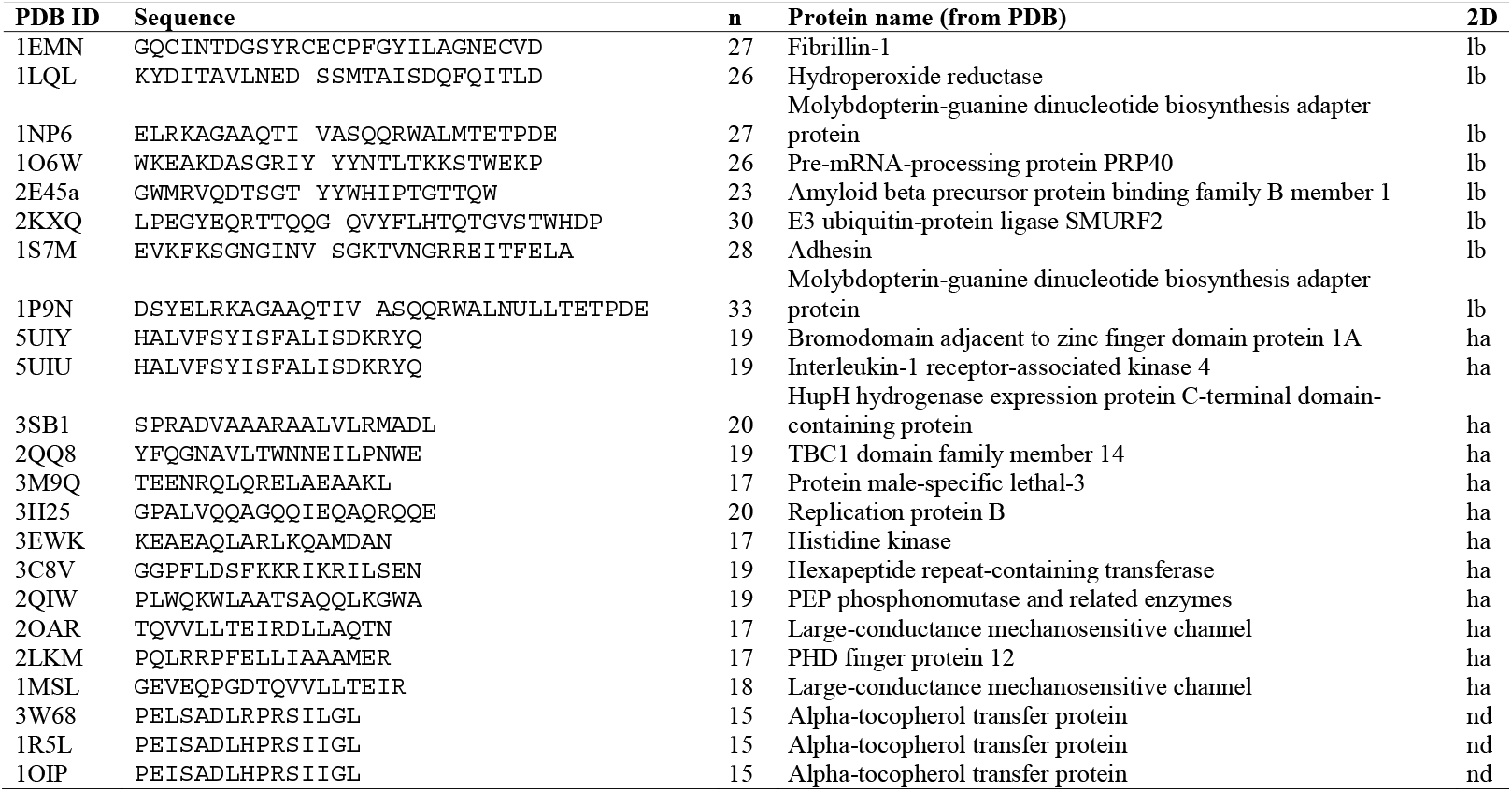
Proteins selected for design using our algorithm. PDB id.: four-letter code of the protein in the PDB, Sequence: the subsequence corresponding to the selected structure, n: length of the selected subsequence, Protein name: the name of the protein in the PDB, 2D: the corresponding secondary structure, ha: alpha helix, lb: beta sheets, nd: not defined.

**Figure S1.**
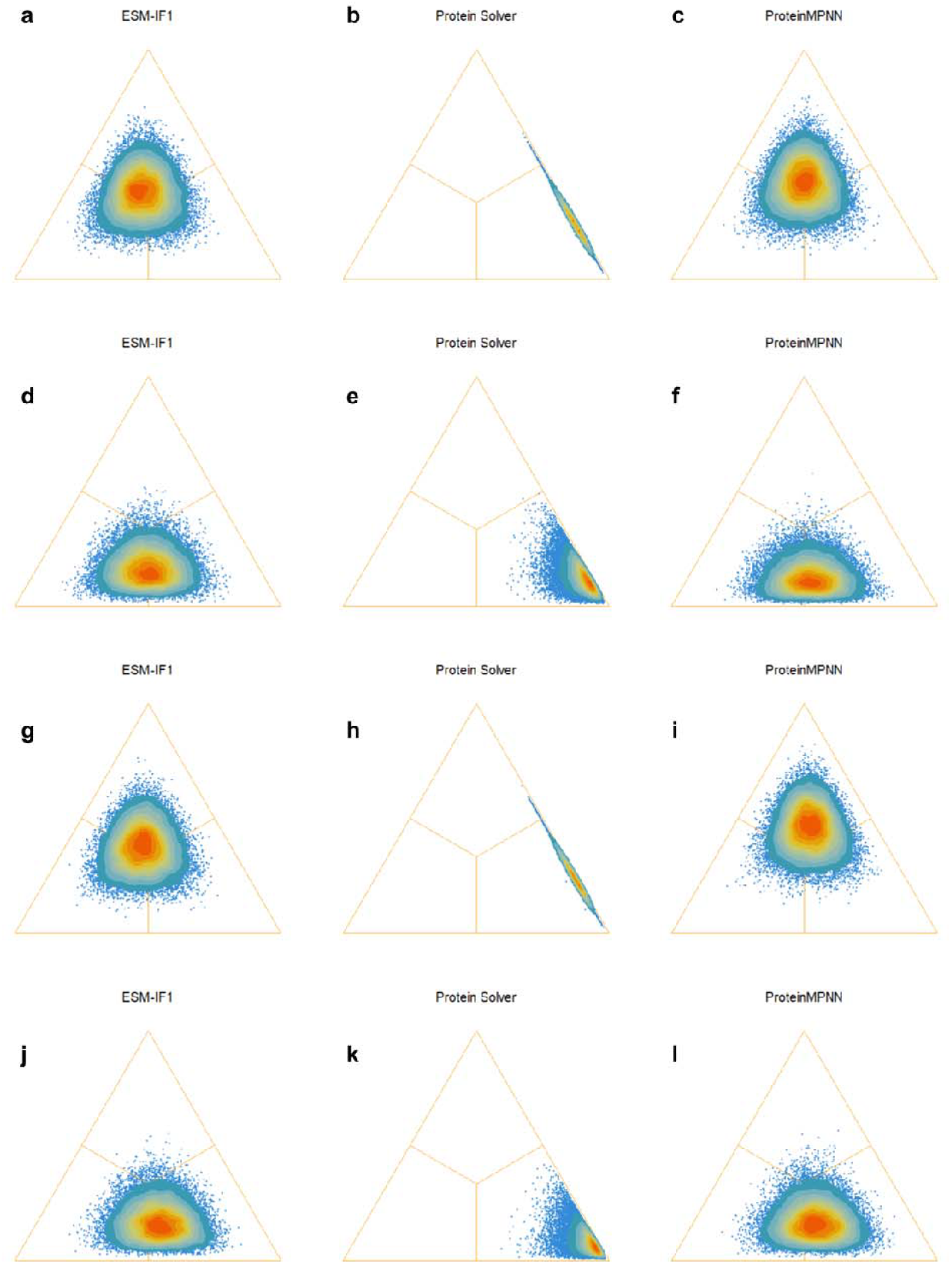
Sign test and Bayesian paradigm, comparison between KCM, ESM-IF1, Protein Solver, and ProteinMPNN. **(a-c)** Comparison of GDT_TS of KCM with respect to **(a)** ESM-IF1, **(b)** Protein Solver, and **(c)** ProteinMPNN, when analyzing the fittest 50 solutions obtained for each protein in each algorithm. **(d-f)** Comparison of RMSD of KCM with respect to **(d)** ESM-IF1, **(e)** Protein Solver, and **(f)** ProteinMPNN, when analyzing the fittest 50 solutions obtained for each protein in each algorithm. **(g-i)** Comparison of GDT_TS of KCM with respect to **(g)** ESM-IF1, **(h)** Protein Solver, and **(i)** ProteinMPNN, when analyzing the fittest 250 solutions obtained for each protein in each algorithm. **(j-l)** Comparison of RMSD of KCM with respect to **(j)** ESM-IF1, **(k)** Protein Solver, and **(l)** ProteinMPNN, when analyzing the fittest 250 solutions obtained for each protein in each algorithm. If the points are concentrated on the left lower corner means that KCM loses, if they are concentrated on the right lower corner then it means that KCM wins. If the points are concentrated on the upper part, both methods perform equally.

**Figure S2.**
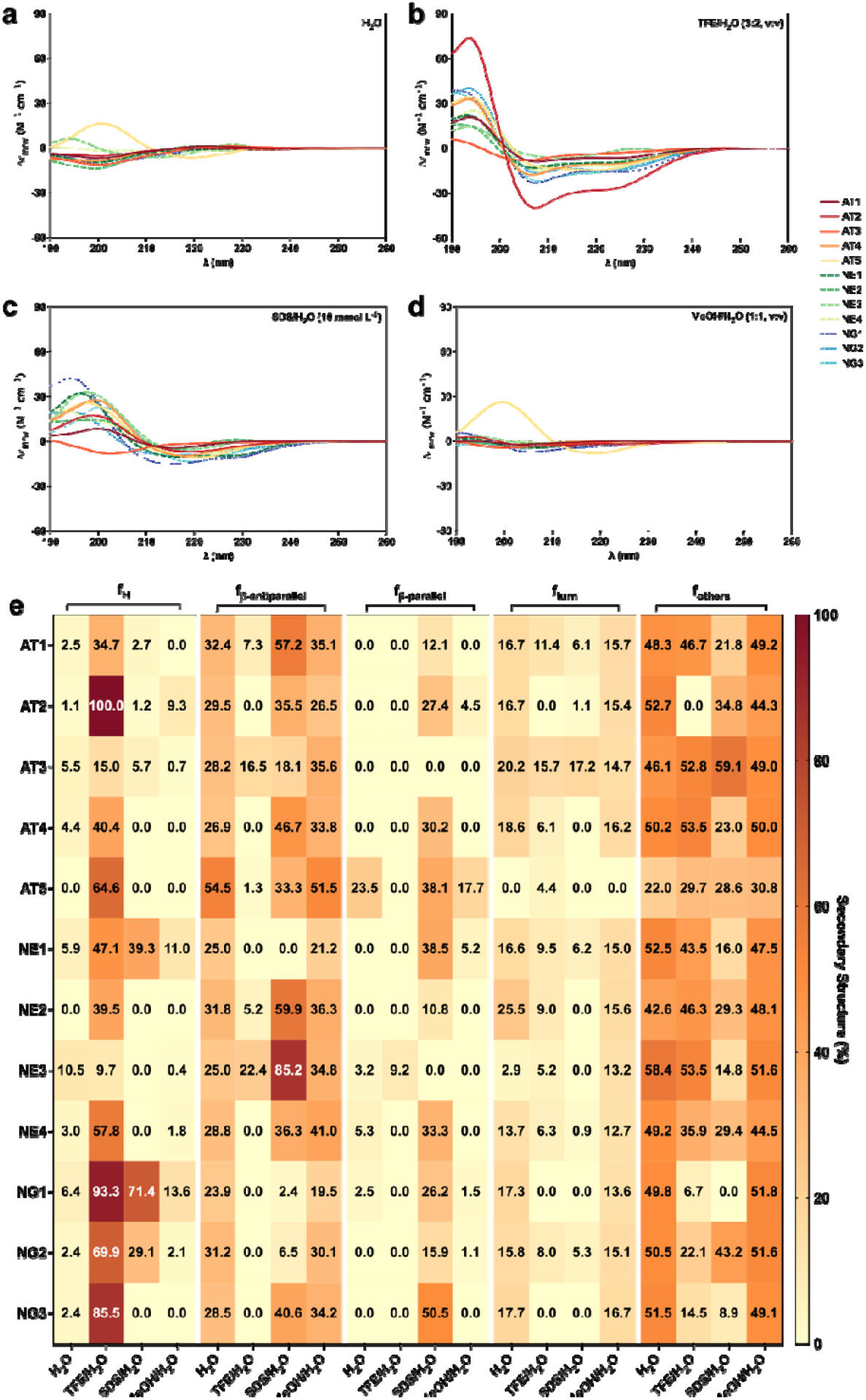
Circular dichroism spectra of designed peptides. Circular dichroism experiments were conducted with the peptides using a J-1500 Jasco circular dichroism spectrophotometer. The spectra were recorded in four different media: **(a)** water, **(b)** 60% trifluoroethanol in water, **(c)** sodium dodecyl sulfate (SDS) in water (10 mmol L^−1^), and **(d)** 50% methanol in water, after three accumulations at 25 °C, using a 1mm path length quartz cell, between 260 and 190 nm at 50 nm min^−1^, with a bandwidth of 0.5 nm. The concentration of all peptides tested was 50 μmol L^−1^. **(e)** Heatmap with the percentage of secondary structure found for each peptide in the four different media analyzed. Secondary structure fraction was calculated using the BeStSel server^7^.

**Figure S3.**
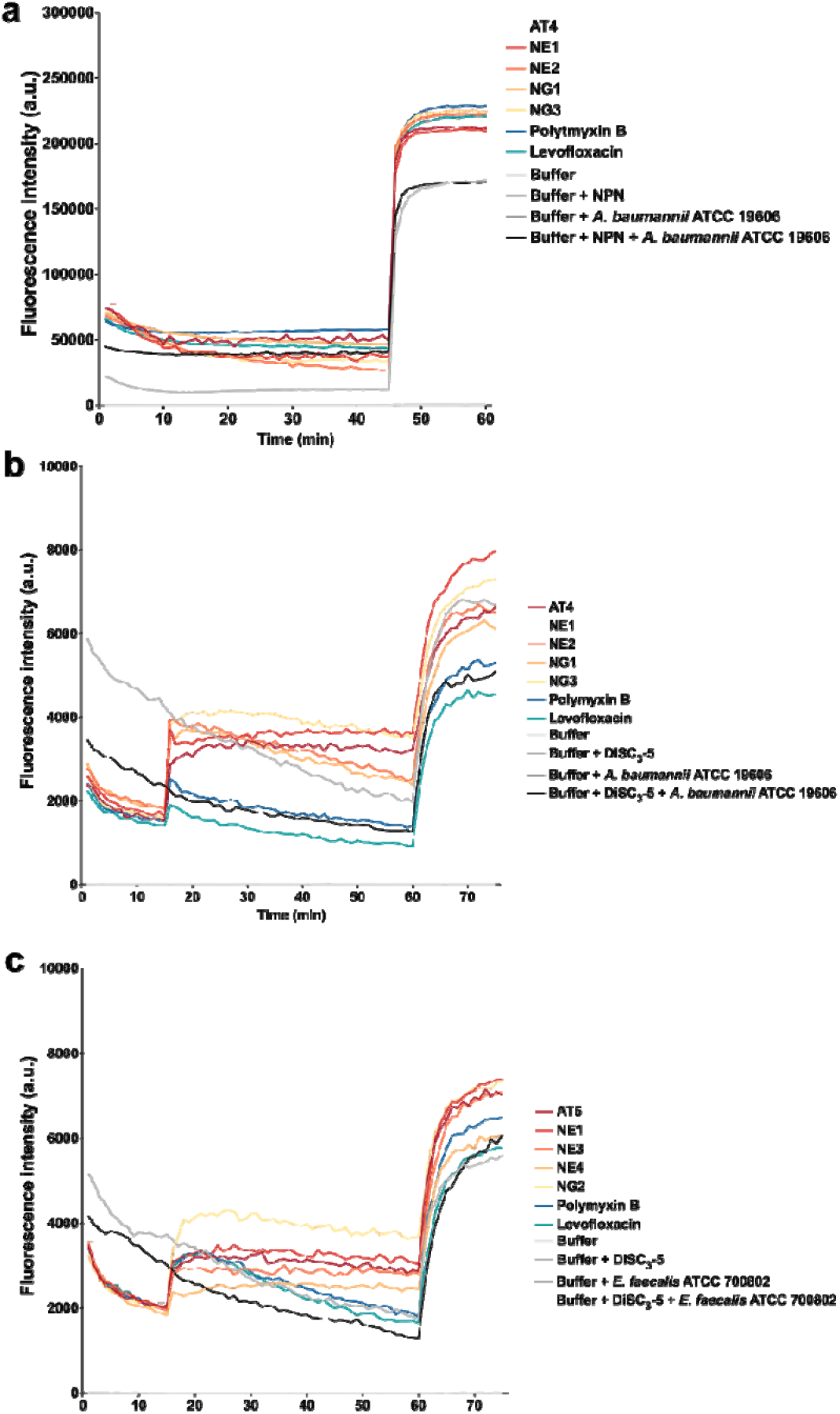
Outer membrane permeabilization and cytoplasmic membrane depolarization of *A. baumannii* ATCC 19606 induced by designed peptides. **(a)** Outer membrane permeabilization was assessed using the probe 1-(N-phenylamino)naphthalene (NPN), showing the permeabilization effects of the designed peptides active against *A. baumannii* ATCC 19606. **(b)** Membrane depolarization assays were performed using the hydrophobic probe 3,3′-dipropylthiadicarbocyanine iodide [DiSC_3_-(5)] on all active peptides against *A. baumannii* ATCC 19606 and vancomycin-resistant *E. faecalis* ATCC 700802. Polymyxin B and levofloxacin served as positive controls, while buffer, buffer with the probe, and buffer with both probe and bacteria were used as baseline controls for fluorescence. The panels display the raw fluorescence intensity data obtained from the experiments. Error bars are the standard deviation obtained from the three replicates.

**Figure S4.**
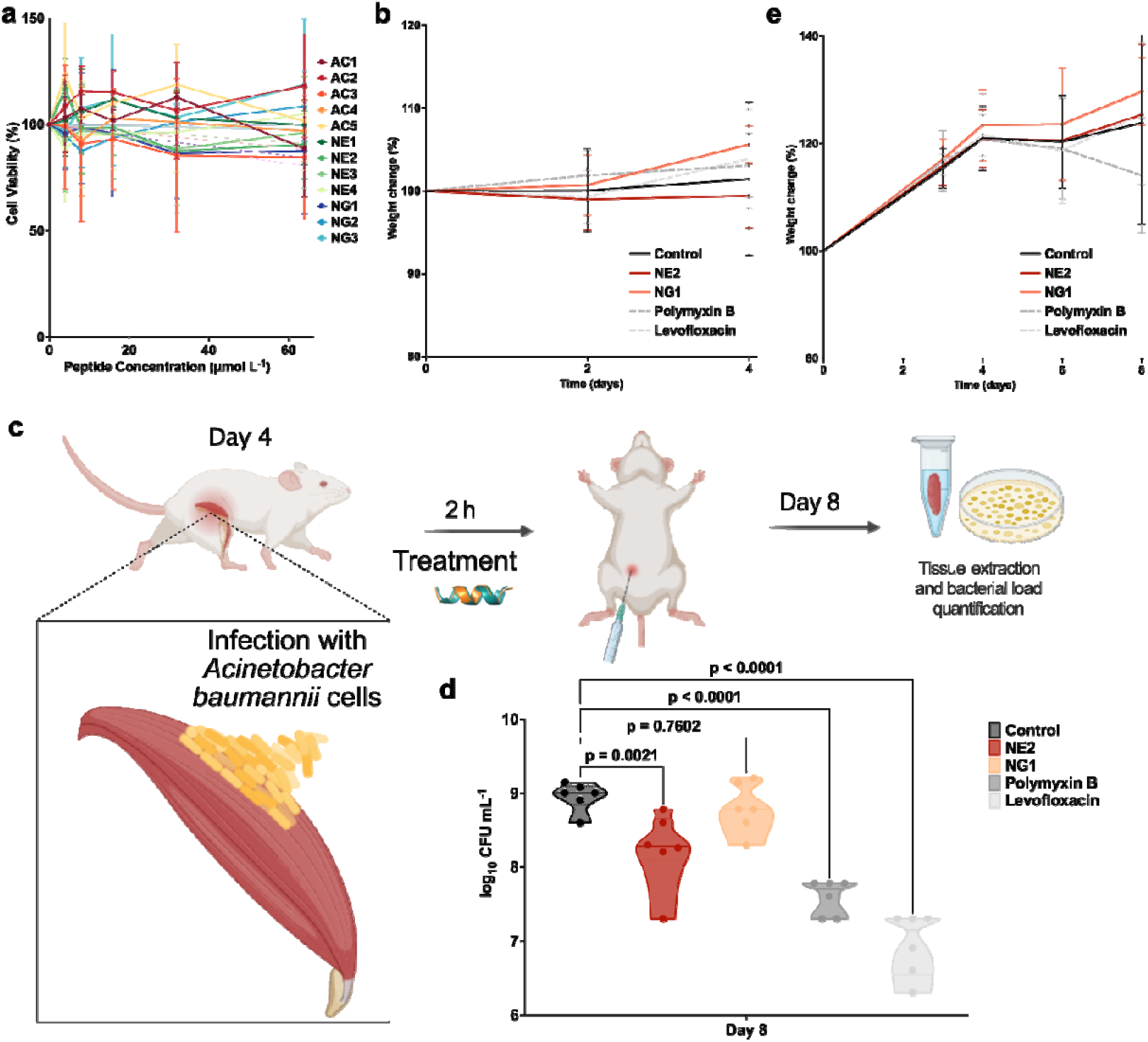
Cytotoxicity and anti-infective activity of designed peptides. **(a)** Cytotoxic effects of increasing concentrations (4, 8, 16, 32, and 34 µmol L^−1^) of each peptide on human embryonic kidney (HEK293T) cells after 24 h of treatment. Mouse weight was monitored throughout the duration of the **(b)** skin abscess model (4 days total) and the **(e)** deep thigh infection model (8 days total) to assess potential toxic effects of both the bacterial load and the designed peptides. **(c)** Schematic of the neutropenic thigh infection mouse model, where the designed peptides were administered intraperitoneally. Anti-infective activity against *A. baumannii* ATCC 19606 was evaluated 4□days after intraperitoneal peptide administration (n□=□6). **(d)** Four days after intraperitoneal injection, NE2 at its MIC (32 μmol L^−1^) inhibited by one order of magnitude the *A. baumannii* ATCC19606 infection, though its activity was less potent than that of polymyxin B and levofloxacin, compared to the untreated control group. Statistical significance in panel **d** was determined using one-way ANOVA followed by Dunnett’s test; P values are shown in the graphs. In the violin, the center line represents the mean, the box limits the first and third quartiles, and the whiskers (minima and maxima) represent 1.5□×□ the interquartile range. The solid line inside each box represents the mean value obtained for each group. Error bars in **a, b** and **e** are the standard deviation obtained from the three biological replicates. Panels **c** was created with BioRender.com.

